# RNA helicase Drs1 gates 25S rRNA domain III incorporation during early nucleolar pre-60S maturation

**DOI:** 10.64898/2026.09.05.749353

**Authors:** Matthias Thoms, Sanem Ayaz-Kök, Kohei Abe, Hussein Hamze, Alina Thielen, Timo Denk, Benjamin Albert, Nika Mikulič Vernik, Leona Chitoiu, Elise Laurent, Ed Hurt, Anthony K. Henras, Roland Beckmann, Valentin Mitterer

## Abstract

Assembly of eukaryotic large ribosomal subunits (LSU) requires coordinated structural and compositional transitions within pre-60S particles, yet the underlying mechanisms remain poorly understood. Here, we show that the DEAD-box helicase Drs1 promotes early maturation across distinct regions of the pre-60S particle. Loss of Drs1 function causes accumulation of co-transcriptional intermediates retaining SSU processome components, indicating that Drs1 promotes timely separation of nascent LSU precursors from the small-subunit assembly pathway. Cryo-EM analyses reveal both a redistribution toward early nucleolar maturation states upon loss of Drs1, including Nsa1-deficient intermediates, and a confinement of Drs1-associated particles to states preceding stable incorporation of 25S rRNA domain III. Drs1 directly engages Erb1 through its unstructured N-terminal extension, promoting stable assembly of the Nop7-Erb1-Ytm1 module associated with domain III maturation. CRAC analysis localizes Drs1 to spatially clustered sites spanning the 5.8S and 25S rRNAs, encompassing domains I–IV. Together, these findings support a model in which Drs1 couples stabilization of assembly-factors with pre-rRNA remodeling across the pre-60S particle, thereby driving ordered early LSU maturation and the timed integration of domain III.

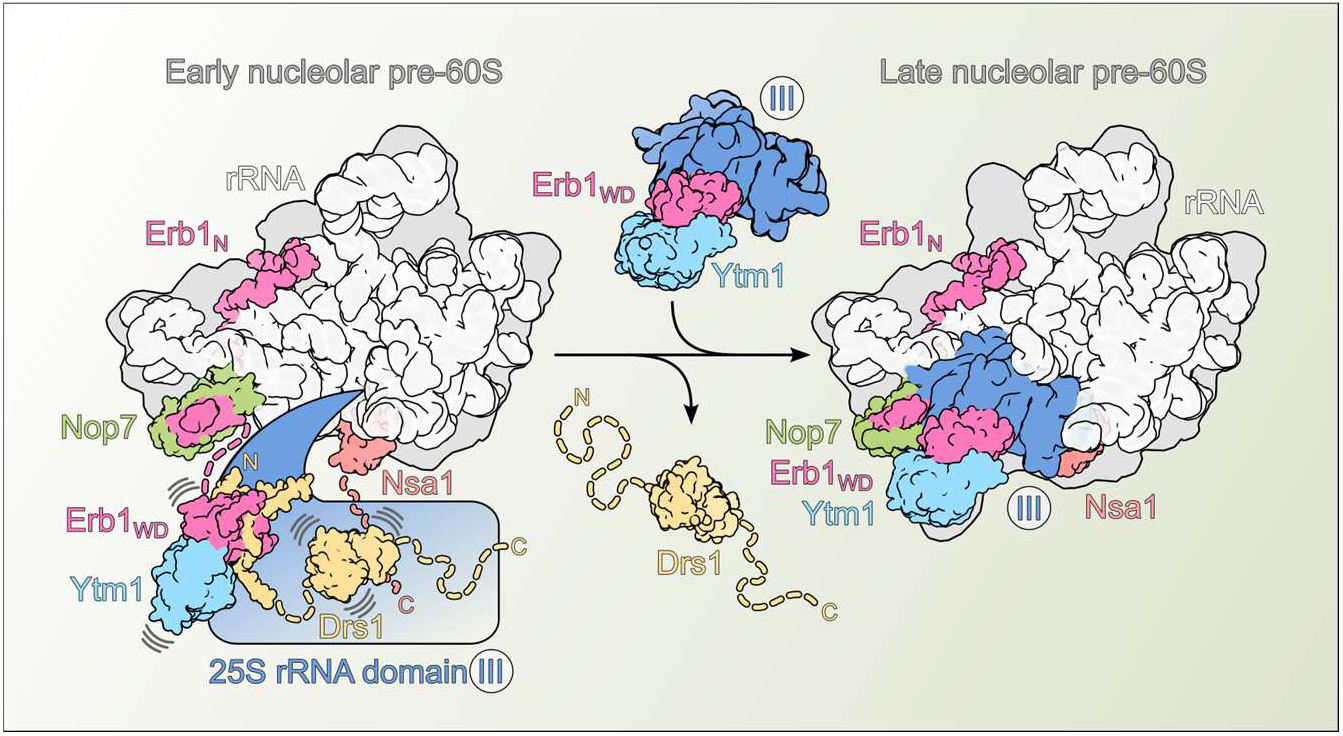

## INTRODUCTION

The synthesis of eukaryotic ribosomes, which are composed of a small (40S) and a large (60S) subunit (SSU and LSU, respectively), is a precisely coordinated and highly dynamic process that spans several cellular compartments, including the nucleolus, nucleoplasm, and cytoplasm. The synthesis pathway depends on the concerted action of over 200 transiently associated protein assembly factors (AFs) and approximately 80 small nucleolar RNPs (snoRNPs), which collectively orchestrate the stepwise formation of functional ribosomal subunits (reviewed in (Baßler and Hurt, 2019; Klinge and Woolford, 2019; Vanden Broeck and Klinge, 2024)).

During ribosome biogenesis, eukaryotic cells must assemble four ribosomal RNA (rRNA) species ─ 18S, 5.8S, 25S (in yeast) or 28S (in humans), and 5S ─ with around 80 ribosomal proteins (r-proteins). The 18S rRNA of the SSU, along with the 5.8S and 25S rRNAs of the LSU, are transcribed as a single precursor RNA (pre-rRNA) by RNA polymerase I. This pre-rRNA harbors external (5’ ETS, 3’ ETS) and internal (ITS1, ITS2) transcribed spacers, which are removed through a series of sequential cleavage and processing events during the maturation pathway (Baßler and Hurt, 2019; de la Cruz et al., 2015; Klinge and Woolford, 2019; Vanden Broeck and Klinge, 2024). The co-transcriptional association of a bulk of assembly factors with the nascent pre-rRNA transcript facilitates the formation of the SSU processome ─ a 6 MDa RNP complex that serves as a structural scaffold, safeguarding the emerging 18S pre-rRNA while coordinating its accurate folding with the integration of SSU r-proteins (Buzovetsky and Klinge, 2025; Chaker-Margot et al., 2017; Dragon et al., 2002; Grandi et al., 2002; Kornprobst et al., 2016; Miller and Beatty, 1969; Singh et al., 2021; Sun et al., 2017). A key maturation event orchestrated within the SSU processome is an endonucleolytic cleavage at site A_2_ within ITS1, which separates the primordial pre-40S and pre-60S precursor particles, thereby committing them to distinct and independent maturation routes (Cheng et al., 2020a; Du et al., 2020; Ismail et al., 2022; Osheim et al., 2004; Wells et al., 2016). However, the timing and precise molecular events leading to this separation remain unclear, including the roles of specific assembly factors. Notably, A_2_ cleavage appears to depend on transcription of the 5.8S and at least part of the 25S rRNA (Khoshnevis et al., 2019; Lebaron et al., 2013; Osheim et al., 2004; Venema and Tollervey, 1996), as well as on the recruitment of early LSU assembly factors, such as Nop4, Nop12, the Npa1 module, and the Rrp5-Noc1-Noc2 complex (Gerhalter et al., 2026; Granneman et al., 2011; Hierlmeier et al., 2013; Ismail et al., 2022; Joret et al., 2018; Rosado et al., 2007; Sanghai et al., 2023; Talkish et al., 2014), which facilitates the formation of a co-transcriptional LSU precursor RNP containing the 5.8S rRNA, ITS2, and 25S rRNA domains I and II (Sanghai et al., 2023). Within this RNP, encapsulation of the 25S rRNA root helix 2 (formed by base pairing between the 5.8S rRNA and the 5’ end of the 25S rRNA) by the Noc1-Noc2 complex establishes a framework for the association of proximal LSU r-proteins and assembly factors like Mak16, Rrp1, and Ebp2. At this stage, the characteristic ITS2-containing foot region of pre-60S particles, along with associated assembly factors including Nop7 and the intertwined N-terminal region of Erb1, is already established (Sanghai et al., 2023).

Following separation from the SSU processome, LSU precursors undergo extensive structural rearrangements driven by a coordinated cycle of assembly factor recruitment and release (Kater et al., 2017; Sanghai et al., 2023; Vanden Broeck and Klinge, 2023; Zhou et al., 2019). During the nucleolar maturation phase, the 5.8S rRNA and the six structural domains of the 25S rRNA (domains I to VI) progressively integrate into the evolving pre-60S core. In the initial assembly intermediates, the 5.8S rRNA, ITS2, and 25S rRNA domains I, II, and VI, adopt a stably folded conformation, forming the solvent-exposed surface of the particle to which the Nsa1 module ─ comprising Nsa1, Mak16, Rpf1, and Rrp1 ─ is assembled (Kater et al., 2017; Sanghai et al., 2018; Vanden Broeck and Klinge, 2023; Zhou et al., 2019). Thereby, the preceding release of Noc1-Noc2 enables the compaction of 25S rRNA domains I and II, as well as the stable integration of domain VI and of assembly factors Nsa1 and Rpf1 (Sanghai et al., 2023). Subsequently, parts of 25S rRNA domain V, followed by domain III and initial segments of domain IV, adopt a folded state before 60S precursors transition to the nucleoplasm (Kater et al., 2020, 2017; Sanghai et al., 2018; Vanden Broeck and Klinge, 2023). To facilitate the nucleolar-nucleoplasmic transition, the Ytm1-Erb1 complex, which serves as a sensor for 25S rRNA domain III maturation proximal to Nop7 at the base of the pre-60S foot (Kater et al., 2017; Sanghai et al., 2018; Thoms et al., 2016), must be released. This critical step is powered by the dynein-related AAA-ATPase Rea1 and its associated pentameric Rix1 subcomplex, inducing large-scale remodeling of the pre-60S subunit as a prerequisite for downstream maturation steps (Bassler et al., 2010; Mitterer et al., 2023).

Prior to the action of the Rea1 AAA-ATPase, several RNA helicases catalyze crucial ATP-driven restructuring steps during nucleolar pre-60S maturation (Aquino et al., 2021; Cruz et al., 2024, 2022; Dembowski et al., 2013; Jaafar et al., 2021; Khreiss et al., 2023; Mitterer et al., 2024, 2023; Portugal-Calisto et al., 2024). Among them is the essential DEAD-box helicase Drs1, which was found on pre-60S particles, including Npa1-, Nsa1-, and Nop7-containing LSU precursors (Dez et al., 2004; Kater et al., 2017; Talkish et al., 2016). Biochemical and genetic data suggest a functional link between Drs1 and the foot factor Nop7 (Adams et al., 2002; Talkish et al., 2016) and, upon shutdown of *de novo* rRNA synthesis, the helicase associates with a ribosome-free Nop7-Erb1-Ytm1 subcomplex (Merl et al., 2010). While cold-sensitive *drs1* mutants, which retain pre-60S binding, exhibit 27SB pre-rRNA accumulation leading to 25S rRNA synthesis defects (Adams et al., 2002; Ripmaster et al., 1992), Drs1 depletion results in the accumulation of the preceding 27SA_2_/A_3_ and 35S pre-rRNAs, accompanied by an enrichment of early pre-60S assembly factors, including the Rrp5-Noc1-Noc2 complex, together with SSU processome components on pre-60S particles (Talkish et al., 2016). This suggests a role of the helicase in early pre-60S maturation at the stage of co-transcriptional ITS1 processing. Like yeast Drs1, its human homolog DDX27 interacts with the PES1-BOP1-WDR12 (Nop7-Erb1-Ytm1) complex while also contributing to 47S (35S in yeast) pre-rRNA processing (Kellner et al., 2015). However, the precise pre-ribosomal binding sites, targets and mechanisms of restructuring, as well as the structural consequences of Drs1 impairment, remain elusive.

Here, we report that the essential DEAD-box helicase Drs1 coordinates a series of critical transitions during early LSU biogenesis. Acute depletion or catalytic inactivation of Drs1 impairs timely separation of nascent pre-60S particles from the SSU processome. At subsequent early nucleolar stages, presence of Drs1 promotes stable association of the solvent-side assembly factor Nsa1. Cryo-EM analyses reveal that loss of Drs1 strongly redistributes pre-60S particles towards early nucleolar intermediates and blocks the progression into pre-60S states with stably assembled 25S rRNA domain III and ES27 of domain IV, whereas a series of high-resolution structures of Drs1-associated particles defines a broad maturation window preceding this transition. We further show that Drs1 stabilizes the Nop7-Erb1-Ytm1 module at the base of the pre-60S foot through a direct interaction of its N-terminal extension with Erb1, while CRAC analysis positions Drs1 at spatially clustered pre-rRNA sites spanning 25S rRNA domains I-IV. Together, our findings support a model in which Drs1 coordinates assembly-factor stabilization and pre-rRNA remodeling across spatially distinct regions of the pre-60S particle, thereby coupling early assembly events around domains I/II to subsequent organization of domain III around the Nop7-Erb1-Ytm1 module.

## RESULTS

### The catalytic activity and unstructured terminal extensions are essential for Drs1 function

To gain deeper insight into the interaction network of the Drs1 helicase (**Figure 1A**), we performed a two-step affinity purification of endogenously N-terminally TAP-Flag (TAPF) tagged Drs1 from yeast whole-cell lysates. Analysis of the final eluate showed that, the helicase efficiently co-purified several nucleolar pre-60S assembly factors, including Ytm1, foot factors Nop7, Rlp7 and Nsa3, as well as components of the Nsa1 module, while very early ribosome assembly factors like the Noc1-Noc2-Rrp5 module did not stoichiometrically co-enrich (**Figure 1B**) (Kater et al., 2017; Sanghai et al., 2018; Zhou et al., 2019). We next fused the endogenous *DRS1* gene to an auxin-inducible degron (AID) tag, allowing rapid degradation of the protein upon addition of the plant hormone auxin (**Figure 1C**). Depletion of Drs1 led to severe growth inhibition on auxin-containing medium, underscoring its essential role in cell viability (**Figure 1D**) (Ripmaster et al., 1992).

**Figure 1.**
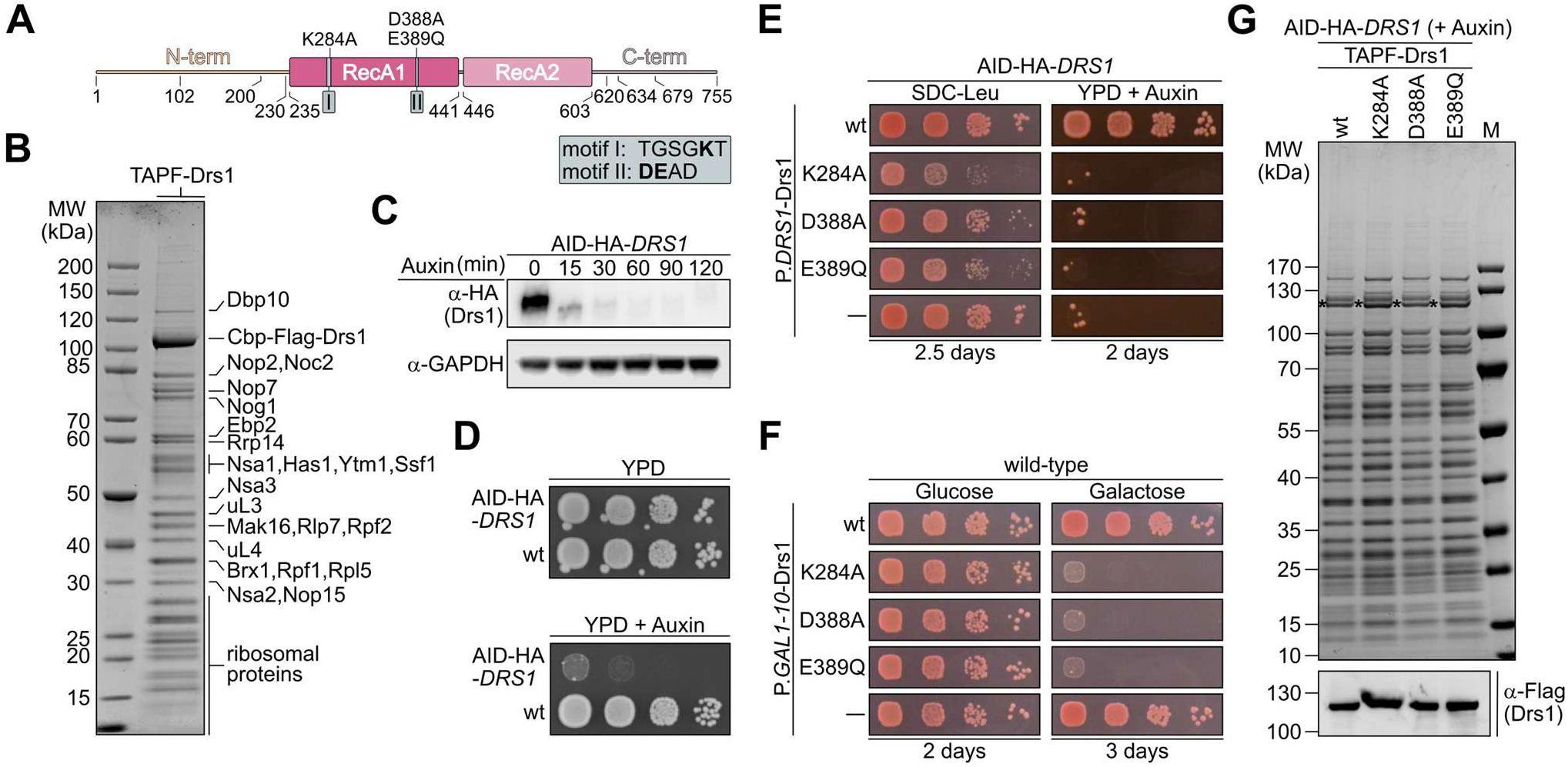
Functional characterization of the pre-60S helicase Drs1. (**A**) Domain organization of Drs1 comprising the catalytic helicase core and unstructured extensions. The conserved motifs I (Walker A) and II (Walker B) required for ATP-binding and hydrolysis, respectively, are depicted. (**B**) Endogenously TAP-Flag (TAPF) tagged Drs1 was tandem-affinity purified and the final Flag eluate was analyzed by SDS-PAGE followed by Coomassie staining. Co-purifying assembly factors are indicated. (**C**) AID-HA-Drs1 is efficiently degraded in the presence of auxin. Logarithmically growing AID-HA-*DRS1* cells were treated with 0.5 mM auxin and the level of AID-HA-Drs1 protein degradation was assessed over 120 min by western blot analysis of whole-cell lysates using an anti-HA antibody. GAPDH served as a loading control. (**D**) The AID-HA-*DRS1* strain was spotted in tenfold serial dilutions on YPD plates without (YPD) or with 2 mM auxin (YPD+Auxin) and incubated for 2 days at 30 °C. (**E**) The AID-HA-*DRS1* strain was transformed with plasmids harboring wild-type *DRS1*, indicated catalytic mutant alleles under the endogenous *DRS1* promoter or empty plasmid (–). Transformants were spotted in tenfold serial dilutions on SDC-Leu (plasmid control) or YPD+Auxin plates and growth was monitored after incubation for 2 days at 30 °C. (**F**) Overexpression of *DRS1* and indicated *drs1* catalytic mutants under the control of the galactose-inducible *GAL1-10* promoter. Transformants were spotted in tenfold serial dilutions on SDC-Leu plates containing glucose (repressed condition) or galactose (induced condition) and growth was monitored after incubation at 30 °C for 2 and 3 days, respectively. (**G**) Tandem-affinity purification of Drs1 wt and mutant constructs. Purifications were performed upon auxin-induced degradation of the AID-tagged wild-type protein, and final eluates were analyzed by SDS-PAGE and Coomassie staining (upper panel) or western blotting (lower panel). The Drs1 bait proteins are marked with an asterisk. M, marker.

Like other DEAD-box proteins, Drs1 contains a catalytic RecA-like core domain that harbors conserved motifs required for ATP binding and hydrolysis, as well as RNA binding (**Figure 1A**; **Supplementary Figure 1A**). To determine how mutations in this catalytic helicase core (Granneman et al., 2006; Ripmaster et al., 1992) affect ribosome assembly, we introduced single-point mutations within the Walker A ATP-binding motif (K284A, motif I) and the Walker B ATP-hydrolysis motif (D388A or E389Q, motif II). Upon expression from plasmids under control of the endogenous *DRS1* promoter, all mutations caused impaired cell growth both upon depletion of endogenous AID-HA-*DRS1* (**Figure 1E**) and in a *DRS1* shuffle strain (**Supplementary Figure 1B**). Moreover, the catalytic mutants exhibited a strong dominant-negative growth phenotype when overexpressed from a galactose-inducible promoter in a wild-type *DRS1* background (**Figure 1F**). Consistent with this dominant-negative effect, TAPF-tagged mutant proteins remained fully capable of binding to pre-ribosomes, as tandem affinity purifications of plasmid-expressed Drs1 variants revealed co-enriched assembly factor patterns comparable to those of wild-type Drs1 particles (**Figure 1G**). Interestingly, deletion of both the unique N- and C-terminal unstructured extensions (**Figure 1A**, **Supplementary Figure 1A**) in the dominant *drs1* K284A mutant abolished its dominant-negative growth phenotype (**Supplementary Figure 1C**, left panel). This effect persisted when the truncation constructs were fused to the SV40 nuclear localization signal (NLS), indicating that the suppression of the phenotype is not due to impaired nuclear import but rather to loss of functional engagement with pre-ribosomes (**Supplementary Figure 1C**, right panel). Together, these findings suggest that catalytically inactive Drs1 effectively binds to pre-ribosomal particles but fails to remodel its substrate and is therefore not efficiently released. The unstructured terminal protein extensions are required for targeting and/or stable engagement of Drs1 with its pre-ribosomal binding sites, thereby enabling its function.

### Drs1 facilitates nucleolar pre-60S assembly during stable incorporation of maturation factor Nsa1

To investigate the function of Drs1 during ribosome assembly, we took advantage of the AID-mediated depletion system and analyzed polysome profiles of AID-Drs1-depleted cells. This revealed a pronounced 60S biogenesis defect after 2 hours of auxin treatment (**Figure 2A**). This defect was evidenced by the accumulation of free 40S subunits, a reduction of both 60S subunits and mature 80S ribosomes, and the appearance of half-mers in the polysome fractions, indicative of mRNA-bound 40S subunits lacking a 60S partner. Western blot analyses of the polysome fractions revealed that the late-joining maturation factor Yvh1 failed to associate with pre-60S particles, shifting instead to low-molecular-weight fractions (**Figure 2A**, right panel). In addition, the early-joining foot-factor Nop7 and the WD40-repeat protein Nsa1, which is positioned at the solvent-exposed surface of nucleolar pre-60S particles, were clearly reduced within the 60S fractions (**Figure 2A**, right panel). These findings demonstrate an effective arrest of 60S synthesis upon Drs1 depletion and indicate that Drs1 promotes an early nucleolar biogenesis step required for the recruitment or stable association of both Nop7 and Nsa1 into pre-60S particles.

**Figure 2.**
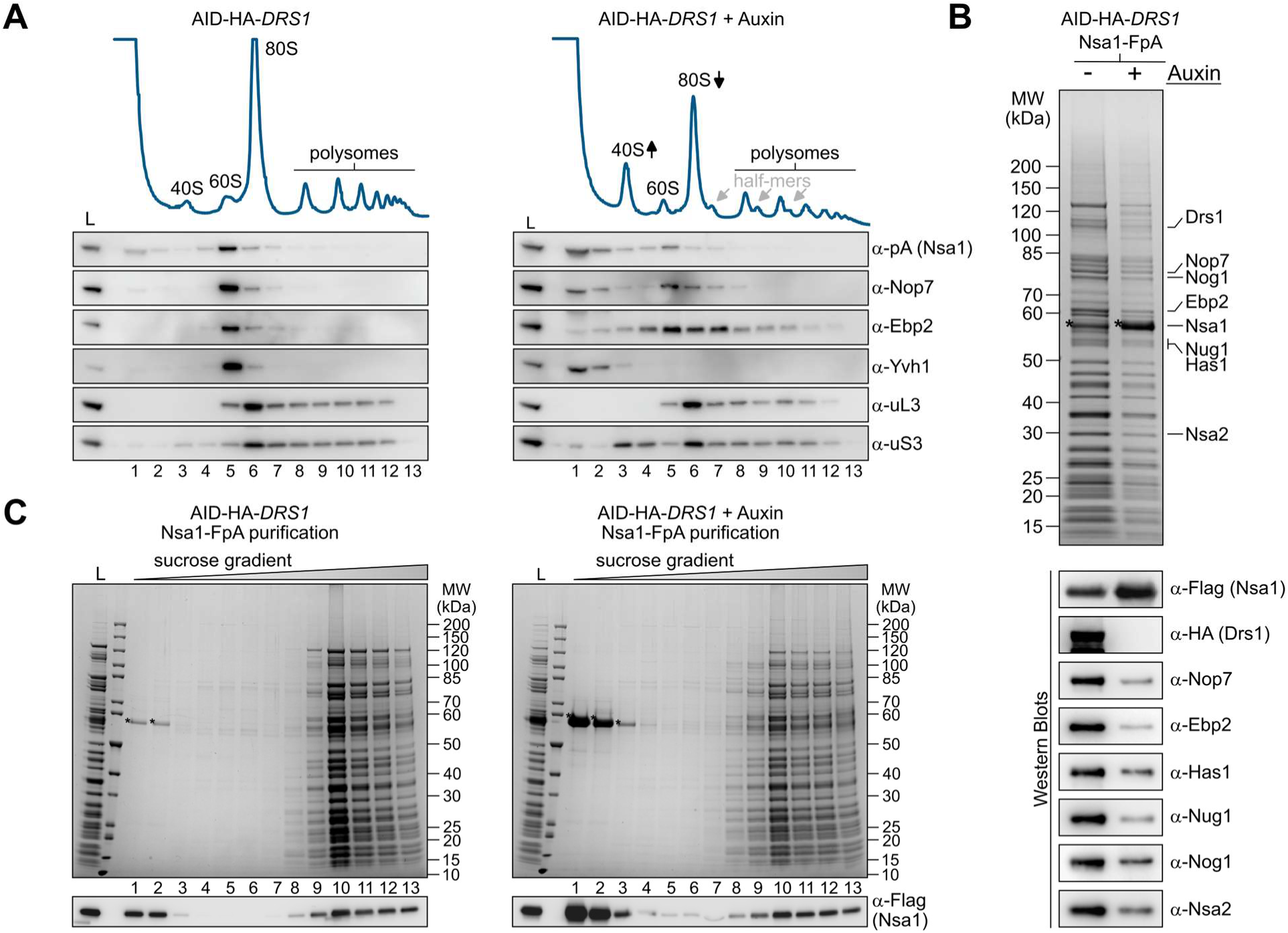
Drs1 promotes pre-60S assembly of Nsa1. (**A**) Polysome profile analysis of AID-HA-*DRS1* cells grown in the absence (left panel) or presence of auxin for 2 h (right panel). Lysates were separated on sucrose gradients and absorbance at 254 nm was recorded. Positions of 40S, 60S, 80S ribosomes, polysomes, and half-mers are indicated. Fractions were collected and analyzed by western blotting using indicated antibodies. L, load. (**B**) Nsa1-FpA was affinity purified from cells in the presence of AID-HA-Drs1 (-Auxin) or upon its auxin-induced degradation (+Auxin). Final eluates were analyzed by SDS-PAGE followed by Coomassie staining (upper panel) or western blotting using the indicated antibodies (lower panels). (**C**) Purified Nsa1-FpA eluates obtained from cells in the presence of AID-HA-Drs1 (left panel) or upon AID-HA-Drs1 depletion (+Auxin) (right panel) were subjected to sucrose gradient centrifugation to separate pre-ribosomal particles from free proteins. Gradient fractions were collected and analyzed by SDS-PAGE followed by Coomassie staining (upper panel) and western blotting using an anti-Flag antibody (lower panel). Fraction numbers increase from low- to high-molecular-weight complexes (left to right). Nsa1-Flag is marked with an asterisk.

Prompted by these observations, we next conducted affinity purifications of Flag-TEV-proteinA (FpA)-tagged Nsa1 from cells either in the presence of auxin (+Auxin; Drs1 depleted) or in its absence (-Auxin; control). Analysis of the obtained Nsa1 eluates revealed a strong reduction of multiple ribosome assembly factors upon Drs1 depletion (**Figure 2B**), indicating impaired Nsa1 incorporation into pre-60S subunits, consistent with our polysome profile data (**Figure 2A**). Further examination of the Nsa1-FpA eluates from Drs1-depleted cells by sucrose gradient centrifugation, which separates pre-ribosomal particles from free proteins and smaller protein complexes, confirmed that a substantial fraction of purified Nsa1 was no longer associated with pre-ribosomes but instead accumulated in the non-assembled low-molecular-weight fractions (**Figure 2C**). Interestingly, a cryo-EM structure of the Noc1-Noc2 RNP (Sanghai et al., 2023), which represents the earliest structurally analyzed co-transcriptional 60S precursor, shows that the maturation factor Noc2 occupies the later Nsa1-Mak16-Rpf1 binding site around ES7A in 25S rRNA domain I on downstream pre-60S intermediates (**Supplementary Figure 2A**-**C**) (Kater et al., 2017; Sanghai et al., 2023). In agreement with a previous study (Talkish et al., 2016), our results therefore suggest that Drs1 depletion may inhibit 60S assembly already at the initial co-transcriptional stage, prior to dissociation of the Rrp5-Noc1-Noc2 module and subsequent stable assembly of Nsa1. Notably, _AlphaFold3_ (Abramson et al., 2024) predicts a direct Drs1-Nsa1 interaction in which the Drs1 RecA2 domain is contacted by the C-terminal α-helix of Nsa1 extending from its WD40 β-propeller (**Supplementary Figure 2D** and **2E**). Whereas the β-propeller is resolved in available pre-60S cryo-EM structures (Kater et al., 2017; Sanghai et al., 2018), the C-terminal helix is not, leaving its orientation on the particles unknown and therefore compatible with the predicted Drs1-binding mode.

### Drs1 promotes the separation of emerging pre-60S particles from the SSU processome

It has been shown previously that Drs1 depletion from a *GAL*-*DRS1* strain leads to accumulation of the 27SA_2_/A_3_ and 35S pre-rRNAs at the whole-cell level, accompanied by persistent association of SSU assembly factors with particles isolated via the pre-60S factor Nop7 (Talkish et al., 2016). Our findings showed that AID-HA-Drs1 depletion weakened the incorporation of both Nop7 and the WD40-repeat protein Nsa1 into pre-60S particles (**Figure 2**). To investigate this phenomenon further, we combined rapid auxin-induced degradation of AID-HA-Drs1 with affinity purification of Nop7-FpA. In contrast to Nsa1 affinity purification, this resulted in a substantial accumulation of very early pre-60S assembly factors (e.g., Noc1, Rrp5, Nop4, Nop12), together with SSU processome factors (e.g., Utp22, Utp20, Utp14, Kre33), on the isolated Nop7 particles (**Figure 3A**). In contrast, factors recruited during later nucleolar and nucleoplasmic pre-60S maturation stages, such as the Spb4 helicase, Nug1, Nog2, Bud20, Arx1 and Rea1, were strongly reduced (**Figure 3A**). Notably, unlike in longer-term GAL-*DRS1* depletion experiments, where SSU, early pre-60S but also later pre-60S factors co-purified (Talkish et al., 2016), immediate Drs1 loss primarily trapped early pre-60S and SSU factors. These results suggest that Drs1 plays a more direct role in promoting A_2_ cleavage and the timely separation of primordial 60S precursors from the SSU processome.

**Figure 3.**
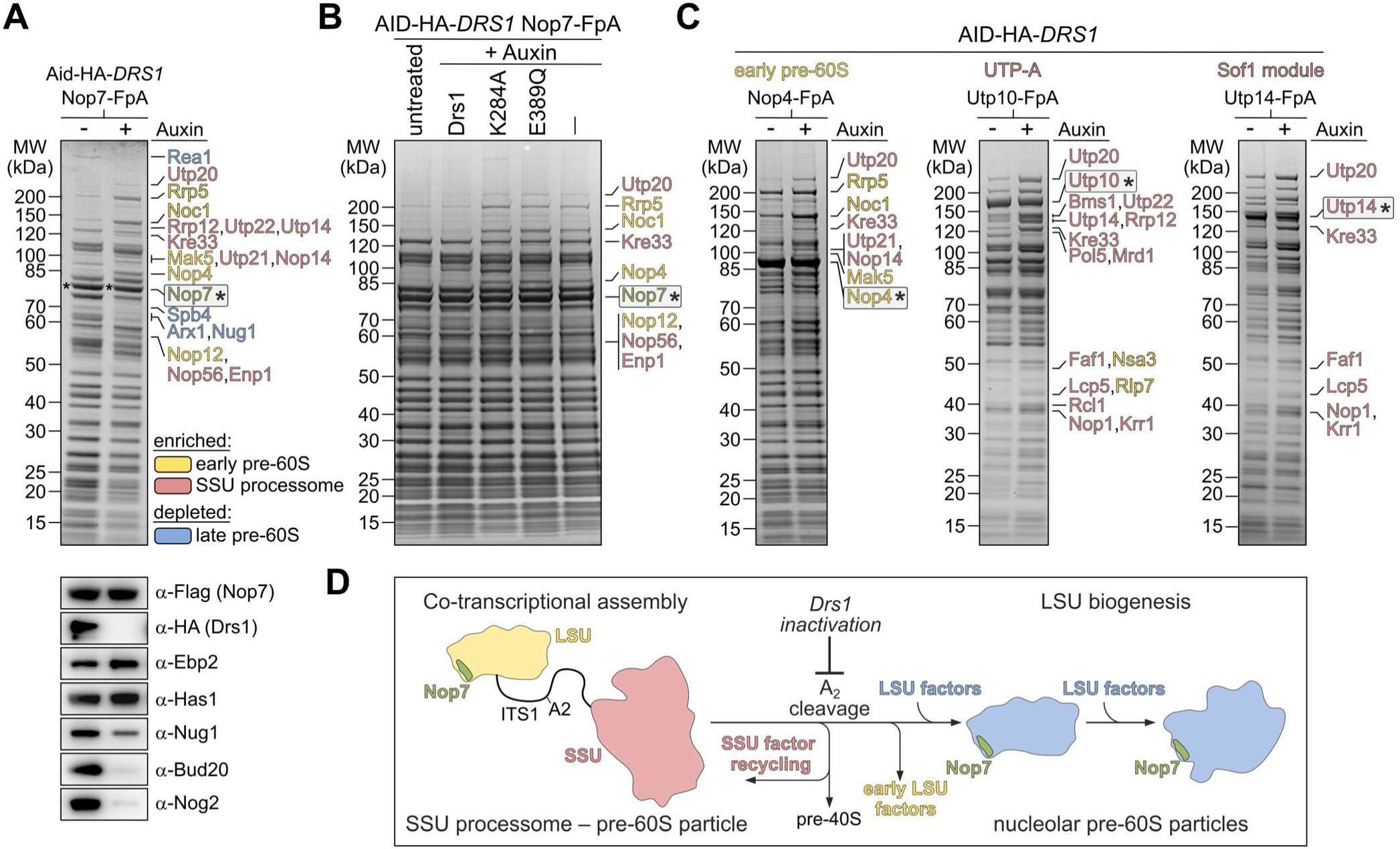
Drs1 promotes separation of pre-60S particles from the SSU processome. (**A**) Nop7-FpA was affinity purified from AID-HA- *DRS1* cells grown in the absence (-Auxin) or presence of auxin (+Auxin). Final eluates were analyzed by SDS-PAGE followed by Coomassie staining (upper panel) and western blotting using the indicated antibodies (lower panel). The Nop7 bait protein is marked with an asterisk. SSU processome factors enriched upon Drs1 depletion are indicated in red, early pre-60S assembly factors increased, and later pre-60S assembly factors decreased upon Drs1-depletion are indicated in yellow and blue, respectively. (**B**) The AID-HA-*DRS1 NOP7*-FpA strain was transformed with plasmids harboring wild-type *DRS1*, the indicated catalytic mutant alleles under control of their native promoter, or empty plasmid (–). Following auxin-induced degradation of endogenous AID-HA-Drs1, Nop7-FpA was affinity purified and final eluates were analyzed by SDS-PAGE followed by Coomassie staining. The Nop7 bait protein is marked with an asterisk. (**C**) Affinity purifications of the Nop4-, Utp10-, and Utp14-FpA bait proteins from AID-HA-*DRS1* cells grown in the absence (-Auxin) or presence of auxin (+Auxin). Final eluates were analyzed by SDS-PAGE followed by Coomassie staining. Asterisks mark the respective bait proteins. SSU processome factors and early pre-60S factors are indicated in red and yellow, respectively. (**D**) Schematic model summarizing the role of Drs1 in co-transcriptional SSU–LSU separation. Drs1 activity enables A2 cleavage, assembly factor recycling, and progression of liberated LSU precursors into nucleolar pre-60S maturation.

We next asked whether this separation requires Drs1’s catalytic activity. Indeed, similar to Drs1-depletion, Nop7 particles purified from *drs1* K284A and E389Q mutant cells also accumulated both early pre-60S and SSU processome factors (**Figure 3B**). Since these *drs1* mutants stably bind to pre-60S particles (**Figure 1G**), we conclude that both the physical presence and the ATP-dependent helicase activity of Drs1 are required for A_2_ cleavage and pre-60S liberation. Supporting this interpretation, the early pre-60S factor Nop4 (Gerhalter et al., 2026; Ismail et al., 2022; Sun and Woolford, 1994) also co-enriched SSU processome components, as well as elevated amounts of Rrp5, Noc1, and Mak5, when isolated from Drs1-depleted cells (**Figure 3C**, left panel). Moreover, affinity purifications via the SSU processome factors Utp10 (Utp-A complex) and Utp14 (Sof1 module) revealed a stronger association of distinct SSU factors such as Utp20, Kre33, and Lcp5 upon Drs1 depletion (**Figure 3C**, middle and right panels). This likely reflects a failure in SSU assembly factor release and recycling due to impaired A_2_ cleavage, reducing the pool of non-assembled Utp10 and Utp14 bait proteins. Consistent with this interpretation, depletion of Drs1 resulted in the accumulation of late SSU processome stages, as indicated by a strong reduction of early-acting assembly factors Pol5 and Mrd1 (**Figure 3C**, middle panel) that are involved in initial Utp-A assembly (Braun et al., 2020; Gallagher, 2019; Gallagher et al., 2004) and 18S rRNA central pseudoknot formation (Lackmann et al., 2018; Segerstolpe et al., 2013), respectively. Together, our data indicate that Drs1’s activity is required to promote the SSU processome-to-pre-60S transition, potentially by mediating RNA restructuring around the A_2_ cleavage site (**Figure 3D**).

### Drs1 depletion traps early assembly intermediates and impairs pre-60S maturation prior to 25S rRNA domain III incorporation

To define the structural consequences of Drs1 depletion and more precisely identify the affected steps within the pre-60S maturation pathway, we next analyzed Nop7-associated particles by single-particle cryo-EM. Nop7-FpA particles were purified from AID-HA-*DRS1* cells grown either in the absence of auxin (-Auxin) or following auxin-induced depletion of Drs1 (+Auxin). The two datasets were collected independently but combined for uniform particle processing and classification, allowing direct and unbiased comparison of the relative representation of individual structural states between the two conditions. After extensive 3D classification, we obtained a series of pre-60S intermediates that could be arranged from early nucleolar to late nucleoplasmic maturation stages in line with previous studies (Hurt et al., 2024; Vanden Broeck and Klinge, 2024) (**Figure 4; Supplementary Figure 3**). In addition to pre-60S particles, we identified SSU processome particles, which were almost exclusively derived from the Drs1-depleted dataset. This observation is consistent with the biochemical accumulation of SSU processome factors on Nop7 particles upon Drs1 depletion and provides independent structural support for delayed A_2_ cleavage and impaired separation of emerging pre-60S particles from the SSU processome. Despite the strong biochemical enrichment of the Rrp5-Noc1-Noc2 factors after Drs1 depletion (**Figure 3A**), we did not obtain reconstructions corresponding to the very early Noc1-Noc2-containing RNPs, potentially reflecting the pronounced aggregation and structural heterogeneity observed upon Drs1 depletion (**Supplementary Figure 3A**, lower panel).

**Figure 4.**
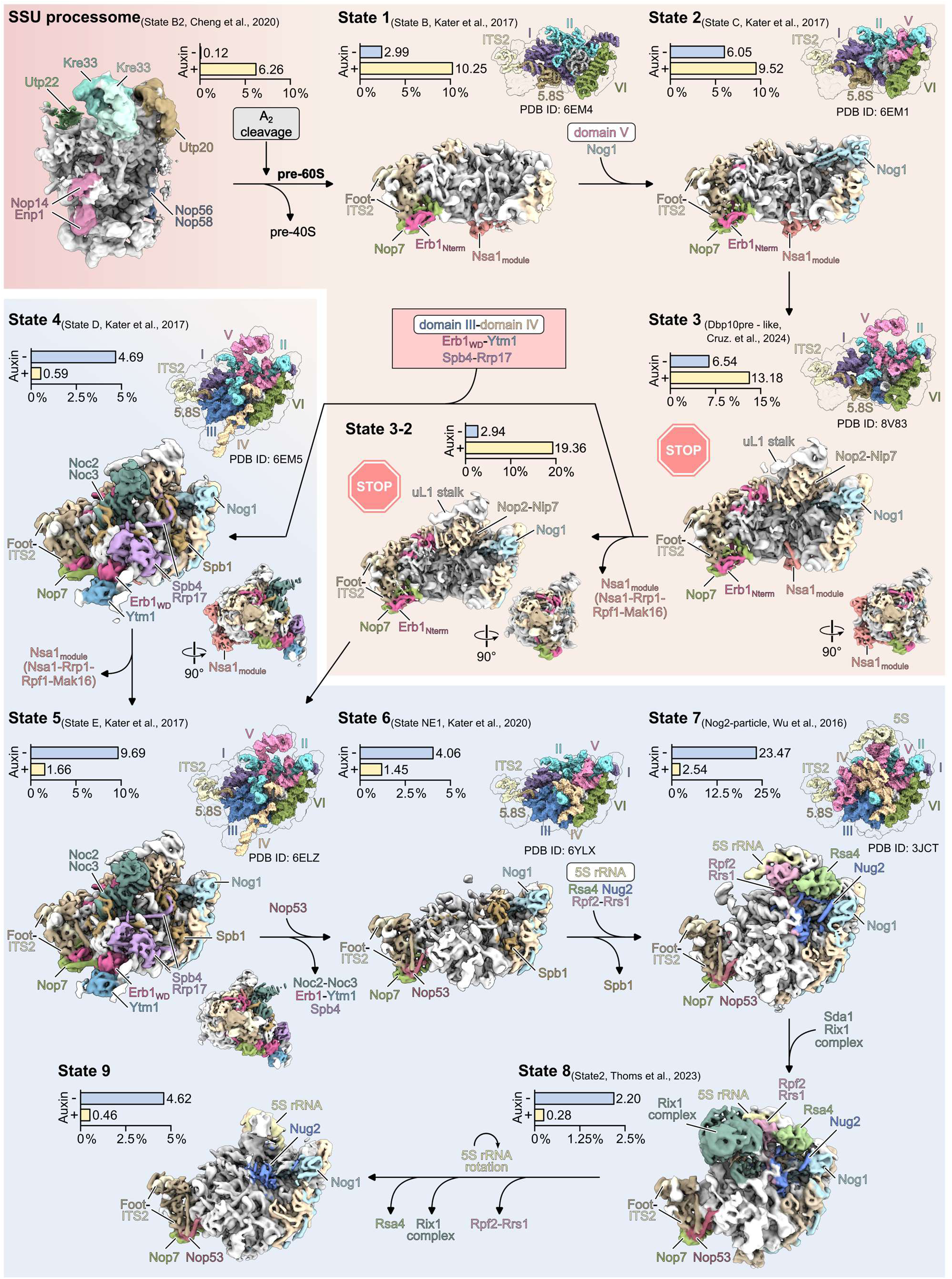
Structural comparison of pre-60S states with and without Drs1. Nop7-FpA was affinity purified from AID-HA-*DRS1* cells grown in the absence (-Auxin, with Drs1) or presence of auxin (+Auxin, without Drs1). The cryo-EM datasets were combined for data processing (see **Supplementary** Figure 3). The cryo-EM densities of the different states are arranged from early to late assembly intermediates, and the maturation trajectory is indicated by arrows. The 25S rRNA domains and AFs that are either assembling on or leaving the particles are indicated above and below, respectively. Individual AFs are colored and labeled, and the remaining AFs are shown in light yellow. The percentages of particles from the parental datasets are shown for each state (upper left). Surface views of matching pre-60S states (PDB IDs are indicated) highlight the assembled rRNA domains of the individual states (upper right) (Cheng et al., 2020b; Cruz et al., 2024; Kater et al., 2020, 2017; Thoms et al., 2023; Wu et al., 2016). The states enriched in the Drs1-depleted dataset (+auxin) and Drs1-containing dataset (-auxin) are highlighted with tan and light blue backgrounds, respectively. Stop signs indicate the block in maturation due to Drs1 depletion.

We next examined how Drs1 depletion affected the distribution of the pre-60S particles themselves. The earliest resolved pre-60S intermediate, State 1, resembles the previously characterized early nucleolar State B (Kater et al., 2017) and contains the 5.8S rRNA together with 25S rRNA domains I, II and VI, whereas the internal domains III–V remain unresolved (**Figure 4**; **Supplementary Figure 3**). The subsequent State 2 additionally contains domain V and the interacting N-terminal domain of Nog1. Further maturation leads to State 3, in which additional elements of domain V, including the uL1 stalk helices and the associated assembly factors Nop2 and Nip7, become stably incorporated. A closely related State 3-2 exhibits an overall similar architecture but lacks the Nsa1 module comprising Nsa1,

Rpf1, Mak16 and Rrp1. Strikingly, all of these early nucleolar intermediates were enriched following Drs1 depletion (**Figure 4**, upper panel). States 3 and 3-2 represented major populations in the Drs1-depleted dataset, accounting for approximately 13.2% and 19.4% of particles, respectively. Compared with the Drs1-containing control, State 3 was enriched approximately 2-fold, whereas State 3-2 showed a much stronger, approximately 6.6-fold enrichment and represented the most abundant pre-60S class after Drs1 depletion. The pronounced accumulation of the Nsa1-deficient State 3-2 is consistent with our biochemical observation of strongly reduced Nsa1 association with pre-60S particles upon Drs1 depletion and suggests premature loss of the Nsa1 module during nucleolar maturation. Notably, in the previously described yeast maturation pathway, release of the Nsa1 module was placed at a later stage, following domain III incorporation and polypeptide exit tunnel (PET) closure, between the previously characterized States D and E (Cruz et al., 2024, 2022; Kater et al., 2017), which correspond to our States 4 and 5 (**Figure 4**). However, related human nucleolar pre-60S intermediates, including states resembling our State 3-2, have been observed both with and without the Nsa1 module (Vanden Broeck and Klinge, 2023), indicating that Nsa1 release is not strictly coupled to domain III/IV incorporation.

A pronounced reversal in particle distribution occurred at all subsequent maturation stages (**Figure 4**, lower panel, **Supplementary Figure 3)**. States 4 and 5 contain stably incorporated 25S rRNA domains III and parts of domain IV, together with the positioned Erb1-Ytm1 module and additional assembly factors including Spb4 and Rrp17, either with (State 4) or without (State 5) Nsa1 module. In contrast to the preceding States 1 to 3/3-2, both intermediates were strongly reduced following Drs1 depletion. This shift towards the Drs1-containing dataset continued throughout the subsequent nucleoplasmic intermediates represented by States 6-9. Most prominently, the well-characterized Nog2-containing State 7 (Leidig et al., 2014; Sekulski et al., 2023; Wu et al., 2016) accounted for approximately 23.5% of particles in the control dataset but only approximately 2.5% of particles following Drs1 depletion. Thus, whereas early nucleolar intermediates accumulate in the absence of Drs1, particles containing stably assembled 25S rRNA domains III and ES27 of domain IV, as well as all subsequently observed maturation states are strongly underrepresented.

Together, the cryo-EM analysis reveals two prominent consequences of Drs1 depletion. First, SSU processome particles accumulate, consistent with delayed A_2_ cleavage and impaired SSU-LSU separation. Second, among particles that have entered the pre-60S maturation pathway, Drs1 loss causes a strong redistribution toward early nucleolar intermediates and markedly impairs progression into states containing stably incorporated 25S rRNA domain III, accompanied by closure of the PET and inter-subunit space. This transition in turn enables stable positioning of the C-terminal Erb1 WD40 β-propeller and its binding partner Ytm1 on the pre-60S particle. Therefore, our structural data identify the transition between States 3/3-2 and State 4/5 (**Figure 4**) as the major Drs1-dependent step in early nucleolar pre-60S maturation.

### Cryo-EM structures of Drs1-associated pre-60S particles

We next sought to define the maturation window during which Drs1 associates with pre-60S particles. To this end, we affinity-purified TAPF-tagged wild-type Drs1 and the catalytically impaired K284A and E389Q variants following depletion of endogenous AID-HA-Drs1 (**Figure 1G**, **5A**) and collected three independent single-particle cryo-EM datasets. Initial processing and classification of each dataset revealed heterogeneous but highly similar populations of pre-60S particles, ranging from early nucleolar State A to C-like intermediates (Kater et al., 2017) to more advanced nucleolar states containing, for example, the stably incorporated uL1 stalk, Nop2-Nip7, and the RNA helicase Dbp10 (**Figure 5B**; **Supplementary Figure 4**). Most importantly, and consistent with our Drs1-depletion analysis (**Figure 4**), all reconstructed particle classes lacked stably incorporated 25S rRNA domain III and therefore represented intermediates preceding the previously characterized States D and E (Kater et al., 2017). Because the three datasets covered a highly similar range of maturation states, we combined them for uniform processing and extensive classification, thereby increasing particle numbers and enabling higher-resolution reconstruction of individual intermediates (**Figure 5B**; **Supplementary Figures 5-9**; **Supplementary Table 1**). The combined analysis resolved a series of closely related nucleolar pre-60S intermediates distinguished by progressive recruitment and rearrangement of 25S rRNA domains, as well as assembly factors and ribosomal proteins, including uL6, Nog1-Nsa2, the uL1 stalk together with Nop2-Nip7, Dbp10, and later nucleolar factors including Noc2-Noc3 and Spb1 (**Figure 5B; Supplementary Figure 10**). Although the datasets were processed together, the origin of particles from the wild-type, K284A and E389Q datasets remained distinguishable, allowing comparison of their relative representation within individual structural classes. Notably, Nsa1-deficient particle classes lacking the Nsa1-Rpf1-Mak16-Rrp1 module were strongly enriched in both catalytic-mutant datasets but only sparsely represented in the wild-type dataset (**Supplementary Figures 4-6**). These particles include the Nsa1-deficient State 3-2 (**Figure 4**) that strongly accumulates upon Drs1 depletion, further indicating that loss of Drs1 activity permits release of the Nsa1 module at this early maturation stage. Despite purification of the particles through Drs1, no additional density corresponding to Drs1 could be identified in any of the reconstructed classes, suggesting that Drs1 is not rigidly positioned onto the pre-60S core, reflecting a flexible mode of association. Together with our Drs1 depletion cryo-EM data, this supports the hypothesis that Drs1 acts on the flexible domain III prior to its stable incorporation.

**Figure 5.**
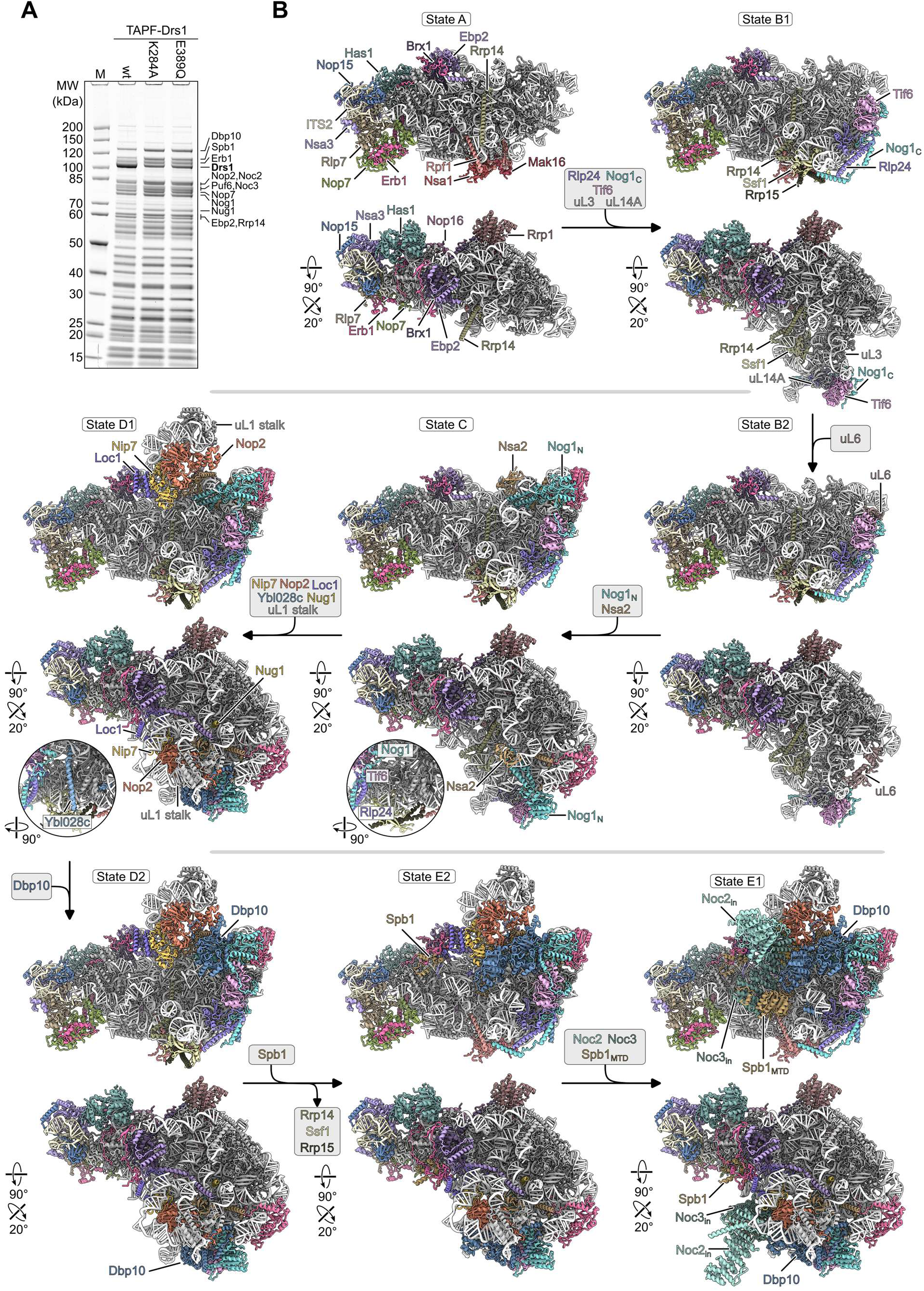
Cryo-EM structures of Drs1-associated early nucleolar pre-60S particles. (**A**) Coomassie stained SDS-PAGE of the Drs1 wt and the catalytic mutants (K284A, E389Q) samples used for single-particle cryo-EM. Plasmids containing TAPF-tagged wild-type *DRS1* or mutant variants under control of the endogenous promoter were purified from the AID-HA-*DRS1* strain after auxin-induced degradation of the AID-tagged wild-type Drs1, and eluates were analyzed by SDS-PAGE and Coomassie staining. Prominent protein bands are labeled. M, marker. (**B**) Molecular models of nucleolar pre-60S states obtained from the combined Drs1 wt/KA/EQ cryo-EM dataset. The intermediates are arranged according to their progression from early toward more advanced nucleolar maturation states. Selected assembly factors that distinguish individual intermediates are colored and labeled. Arrows indicate the maturation trajectory. Magnified views of States C and D1 highlight the region around Nog1, Tif6, and Rlp24 and the recruitment of Ybl028c in State D1.

Several of the resolved structures correspond to previously characterized yeast pre-60S States A–C (Kater et al., 2017). However, the high-resolution reconstructions, reaching approximately 2.3–2.7 Å for several of these intermediates, enabled more detailed structural models than the previously available ones (**Supplementary Figures 7–9**; **Supplementary Table 1**). Beyond these previously characterized states, we observed intermediates that have not been structurally characterized in *S. cerevisiae* but closely resemble related states described in human (Vanden Broeck and Klinge, 2023) and *Chaetomium thermophilum* (Lau et al., 2023), revealing a highly conserved sequence of structural transitions during early eukaryotic LSU assembly (**Figure 5**; **Supplementary Figures 5– 9**).

### Drs1 directly interacts with Erb1 and orchestrates timely 25S rRNA domain III integration

All resolved Drs1-associated intermediates preceded stable incorporation of 25S rRNA domain III (**Figure 5B**). To visualize the structural transition following the Drs1-associated maturation window, we compared Drs1-associated State E1 with a post-Drs1 pre-60S state (**Figure 6**) (Kater et al., 2017; Mitterer et al., 2023). During this transition, stable incorporation of domain III facilitates PET closure, while the C-terminal Erb1 WD40 β-propeller together with Ytm1 and Spb4 become stably positioned around the integrated ES27 domain IV (**Figure 6B**-**C**; **Supplementary Figure 10**). Together with previous biochemical and genetic studies linking Drs1 to the Nop7-Erb1-Ytm1 module (Adams et al., 2002; Merl et al., 2010; Thoms et al., 2016), these structural observations raised the possibility that Drs1 directly contributes to its stabilization during this transition. To investigate this, we analyzed Nop7-FpA particles following Drs1 depletion by sucrose gradient centrifugation (**Figure 7A**).

**Figure 6.**
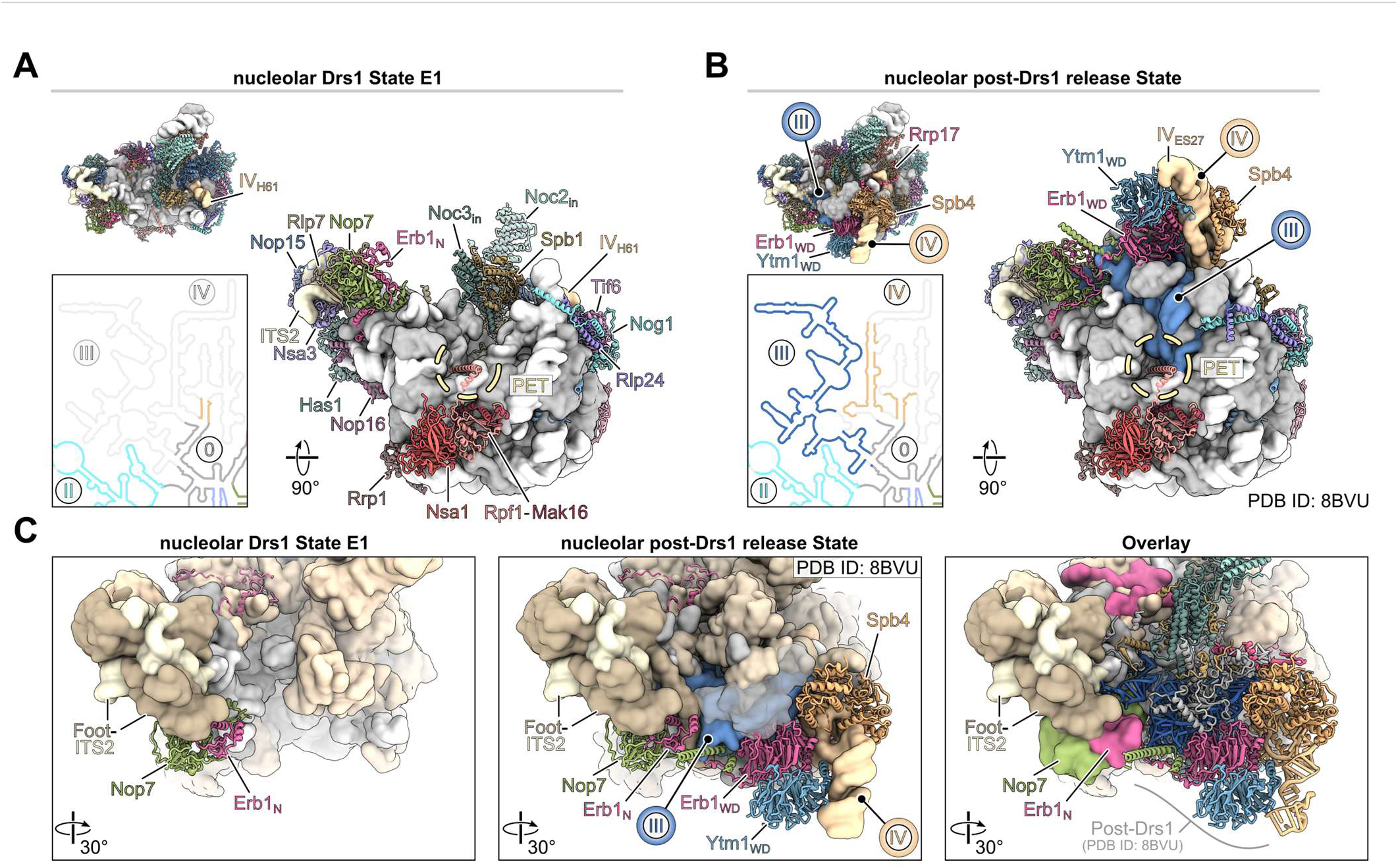
Structural comparison of Drs1-bound and post-Drs1 pre-60S States. (**A**-**B**) Comparison of the Drs1-bound pre-60S State E1 (A) and a subsequent nucleolar pre-60S State after Drs1 release (B) (PDB ID: 8BVU) (Kater et al., 2017; Mitterer et al., 2023). The pre-60S states are shown from the intersubunit side (upper left) and the PET region (right panels). Ribosomal proteins and rRNA are shown as filtered surface representations in gray and light gray, respectively. The 25S rRNA domains III and IV embedded in the post-Drs1 State are highlighted in blue and orange, respectively (B). Associated AFs are presented as colored ribbon models. AFs present in both states are labeled in (A), and AFs that associate with the post-Drs1 release state are labeled in (B). Close-ups of the mature 25S rRNA secondary structure focusing on domains III and IV are shown, and rRNA regions presented in the models are colored. Flexible regions are shown in transparent gray (lower left panels). (**C**) Close-up views of the Drs1 State E1 (left panel) and the post-Drs1 State (middle panel), focusing on the ITS2-containing Foot region and the binding region of 25S rRNA domains III and IV. For clarity, the pre-60S states are shown as filtered surface representations, with AFs in light orange, rRNA and r-proteins in gray, and Foot AFs and ITS2 in brown and yellow, respectively. Relevant AFs, including Nop7-Erb1-Ytm1, are shown as ribbon models, and 25S rRNA domains III and IV are colored and highlighted. The overlay (right panel) shows the rigid body fit of Drs1 State E1 and the post-Drs1 State. For better visualization, the Drs1 State E1 is displayed as surface representation and the overlayed post-Drs1 State as ribbon model, highlighting the incorporation of domain III/IV and associated AFs and r-proteins after Drs1 release.

**Figure 7.**
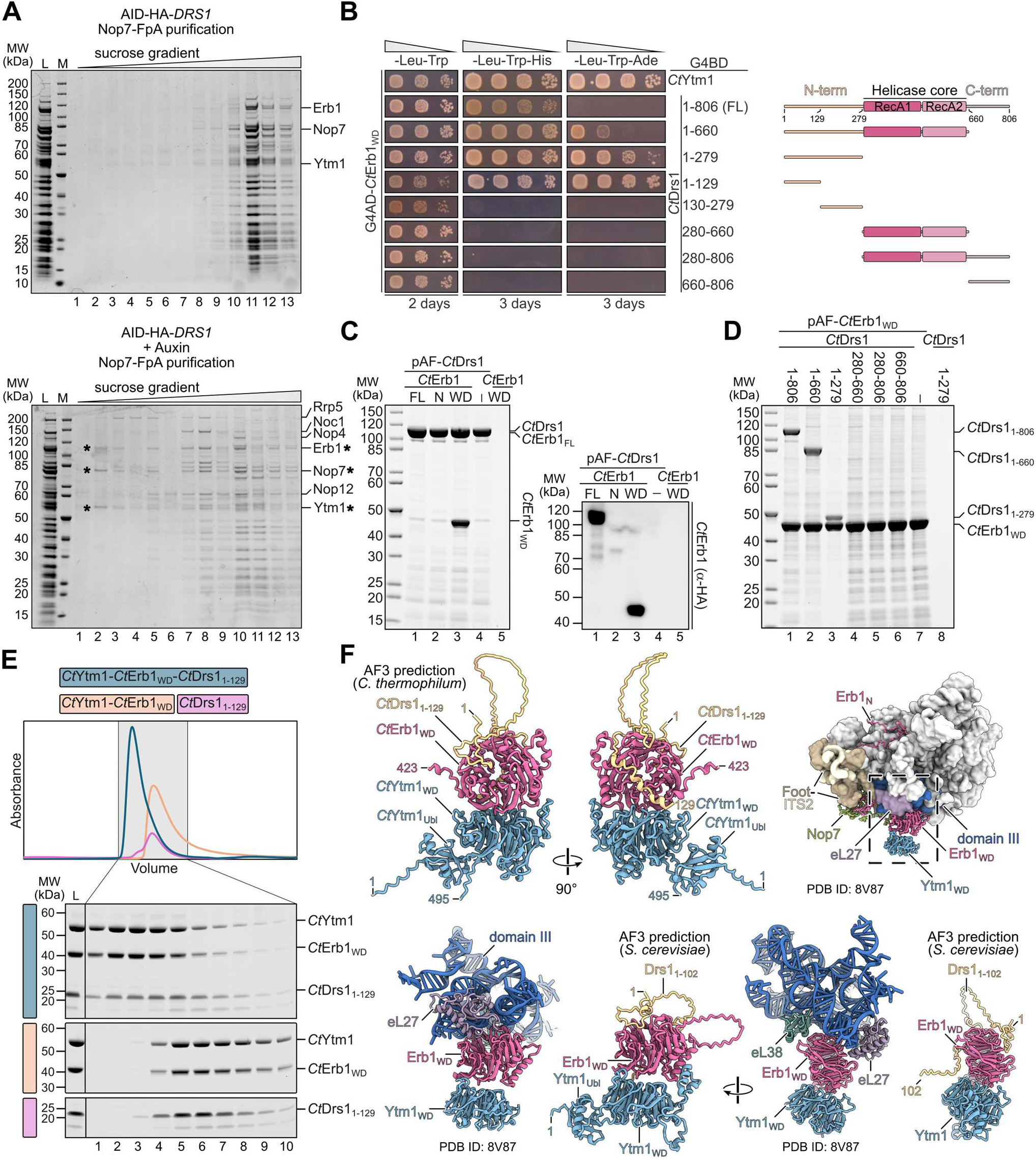
Drs1 promotes stable assembly of the Nop7–Erb1–Ytm1 module and associates with Erb1. (**A**) Redistribution of Nop7-associated particles upon Drs1 depletion. Nop7-FpA eluates purified from AID-HA-*DRS1* cells in the presence (upper panel) or absence (lower panel) of AID-HA-Drs1 were subjected to sucrose gradient centrifugation. Gradient fractions were collected and analyzed by SDS-PAGE and Coomassie staining. Relevant early pre-60S assembly factors and the Nop7–Erb1–Ytm1 module are indicated (*). M, Marker; L, Load (**B**) Yeast two-hybrid interaction between Drs1 and Erb1 constructs from *C. thermophilum* (*Ct*). Indicated *Ct*Drs1 constructs and *Ct*Ytm1 fused to the Gal4 DNA-binding domain (G4BD) were tested for interaction with the C-terminal *Ct*Erb1-WD40 (WD) domain fused to the Gal4 activator domain (G4AD). Cells were spotted in tenfold serial dilutions on SDC-Leu-Trp (plasmid control), SDC-Leu-Trp-His (growth indicates weak interaction), and SDC-Leu-Trp-Ade (growth indicates strong interaction) plates and incubated at 30 °C for the indicated times. (**C**-**D**) The N-terminus of *Ct*Drs1 physically interacts with the C-terminal *Ct*Erb1 WD40 domain. Indicated Drs1 and Erb1 constructs under the control of the *GAL1-10* overexpression promoter were co-expressed in yeast. (**C**) Full-length pA-TEV-Flag-*Ct*Drs1 (pAF) was purified from yeast cells expressing the indicated 2xHA-tagged *Ct*Erb1 constructs. Final elutes were analyzed by SDS-PAGE and Coomassie staining. (**D**) The pA-TEV-Flag-tagged *Ct*Erb1WD construct (aa 423-802) purified from yeast cells co-overexpressing the indicated *Ct*Drs1 constructs. Eluted proteins were analyzed by SDS-PAGE and Coomassie staining. (**E**) Size-exclusion chromatography of the reconstituted *Ct*Ytm1–*Ct*Erb1WD–*Ct*Drs11–129 complex, the *Ct*Ytm1–*Ct*Erb1WD, and the *Ct*Drs11–129 fragment alone. Elution profiles are shown above, and corresponding fractions were analyzed by SDS-PAGE followed by Coomassie staining. L, Load. (**F**) AlphaFold3 predictions of the *Ct*Ytm1–*Ct*Erb1WD in complex with the *Ct*Drs11–129 N-terminus from *C. thermophilum* (upper left panels) and positioning of the yeast Nop7–Erb1–Ytm1 subcomplex within the cryo-EM model of a nucleolar pre-60S (Dbp10 post-catalytic, PDB ID: 8V87) after 25S rRNA domain III incorporation and Erb1WD-Ytm1WD binding (upper right). Magnified views of the Erb1WD-Ytm1WD in contact with 25S rRNA domain III and ribosomal proteins eL27 and eL38 within the pre-60S particle (PDB ID: 8V87) compared to AlphaFold3 predictions of the Ytm1–Erb1WD–Drs11–102 complex from *S. cerevisiae* shown in the same orientations (lower panels).

Interestingly, in the absence of Drs1, Nop7-associated factors were clearly redistributed across the gradient, with distinct smaller RNP complexes containing early factors Rrp5, Noc1, Nop4, and Nop12 migrating to lower molecular-weight fractions (**Figure 7A**, lower panel ─ lanes 5, 7, 8) relative to the major pre-ribosomal fractions of wild-type eluates (**Figure 7A**, upper panel — lanes 11, 12, 13). Most notably, the Nop7-Erb1-Ytm1 subcomplex (Miles et al., 2005; Tang et al., 2008; Thoms et al., 2016), which associates with the pre-60S foot and contacts 25S rRNA domain III as well as the base of ES27 in domain IV (**Figure 6B-C**), was partially released, migrating in the top fractions of the gradient (**Figure 7A**, lower panel, lane 2), indicating its destabilization and detachment from pre-ribosomal particles upon Drs1 loss. We conclude that, while initial docking of the Nop7-Erb1-Ytm1 module can occur without Drs1, the helicase facilitates its final integration into pre-60S particles, a step that could be coupled to compaction and folding of the adjacent 25S rRNA domain III during Drs1 release.

To explore whether Drs1 directly engages the Nop7-Erb1-Ytm1 module, we performed yeast two-hybrid interaction assays using constructs from the thermophilic fungus *Chaetomium thermophilum*, which yielded more robust results in this assay than the equivalent yeast constructs (**Supplementary Figures 1A** and **11A**–**C**). Indeed, full-length *Ct*Drs1 interacted with the *Ct*Erb1 WD40 C-terminal domain but not the *Ct*Erb1 N-terminal domain, and the interaction was even stronger with an N-terminal Drs1 fragment (aa 1-279) (**Figure 7B**; **Supplementary Figure 11A**–**C**). Further truncation analyses showed that, consistent with human studies (Kellner et al., 2015), the first half of the unstructured N-terminal extension of Drs1 (aa 1-129) was sufficient for the interaction with Erb1, whereas the second half (aa 130-279) showed no interaction (**Figure 7B**, **Supplementary Figure 1A**). In agreement, the catalytic core (aa 280-660) and the C-terminal helicase extension (aa 660-806) did not mediate interaction. To corroborate these results, we co-overexpressed and purified pAF-tagged *Ct*Drs1 along with 2xHA tagged *Ct*Erb1 truncation constructs. The Drs1 purifications co-enriched full-length *Ct*Erb1, as well as the C-terminal WD domain of *Ct*Erb1 but failed to interact with the N-domain of *Ct*Erb1 (**Figure 7C**). Consistently, purification via the pAF-tagged C-terminal domain (WD) of *Ct*Erb1 together with *Ct*Drs1 truncations confirmed direct interactions with Drs1 constructs harboring the N-terminal region including the full length Drs1 (aa 1-806), a truncation lacking the C-terminal extension (aa 1-660) and the N-terminus of Drs1 (aa 1-279) itself (**Figure 7D**, lanes 1-3). In agreement, truncations lacking the Drs1 N-terminus all failed to interact with the Erb1 WD40 β-propeller. (**Figure 7D**, lanes 4-6; **Supplementary Figure 11D**). Moreover, size-exclusion chromatography after *in vitro* reconstitution demonstrated that *Ct*Ytm1, *Ct*Erb1, and the first half of the unstructured *Ct*Drs1 N-domain (aa 1-129) form a stable trimeric complex that co-eluted as a single peak, indicating robust association (**Figure 7E**). Finally, AlphaFold3 multimer predictions (Abramson et al., 2024) provided a structural framework for the experimentally defined interaction. The predicted trimeric complex shows that the Drs1 N-terminal region and Ytm1 can simultaneously engage the C-terminal Erb1 WD40 β-propeller, consistent with the stable *Ct*Ytm1–*Ct*Erb1WD–*Ct*Drs1(1–129) complex observed by size-exclusion chromatography (**Figure 7E**-**F**; **Supplementary Figure 12**). Comparison with nucleolar pre-60S structures further places the Drs1 N-terminal region adjacent to the site at which the 25S rRNA domain III subsequently engages Erb1 (**Figure 7F**). This arrangement is consistent with a gating mechanism in which Drs1 transiently occupies the domain III assembly site, thereby blocking its engagement with the Erb1 WD40 β-propeller until Drs1-dependent remodeling events have been completed. Together, these data support a model in which Drs1 stabilizes the Nop7-Erb1-Ytm1 module within early pre-60S particles through a direct N-terminal contact with Erb1 while coordinating the timing of subsequent domain III incorporation.

### Drs1 crosslinks to spatially clustered sites across the 5.8S and 25S pre-rRNA

To determine the rRNA binding sites of Drs1, we performed *in vivo* UV <u>cr</u>osslinking and <u>a</u>nalysis of <u>c</u>DNA (CRAC) (Granneman et al., 2009) using cells expressing Drs1 tagged at its N-terminus with proteinA-TEV-(His)_6_ (pATH). Sequencing of the cDNA library revealed several prominent hit peaks across the 5.8S and 25S rRNAs, but not in the 18S rRNA sequence or spacer regions (**Figure 8A**-**B**). Drs1 crosslinked to the central part of the 5.8S rRNA (nt 33-138), as well as to multiple regions within the 25S rRNA, including domain I (nt 10-64, 322-378), domain II (nt 881-928), domain III (nt 1452-1477, 1601-1619 and 1838-1879) and domain IV (nt 1911-1937, 2091-2120) as well as in neighboring regions within the root helices (nt 1448-1451, 1880-1909, and 1909-1910) (**Figure 8A**-**B**). Although the CRAC signals were broadly distributed over the linear sequence of 5.8S and 25S rRNAs, their mapping onto pre-60S structures revealed that they cluster within a spatially confined region of the pre-rRNA (**Figure 8C**-**D**). The cross-linking sites in domains III and IV correspond to rRNA areas which remain unfolded in early nucleolar Drs1-associated states and become progressively folded during subsequent maturation towards states D/E (**Figure 6**) (Cruz et al., 2024; Kater et al., 2017). In contrast, the binding sites within the 5.8S and 25S rRNA domains I and II are located in the assembled core of the particle. However, they are found at the periphery of the not-yet-incorporated domain III, in direct proximity to Nop7 and the N-terminal region of Erb1, both of which interact with domain III (**Figure 8C**-**D**; lower panels). Taken together, the CRAC data position Drs1 within a spatially confined pre-rRNA region encompassing domains I-IV and is consistent with its extended domain architecture, allowing contacts with distinct rRNA elements during early nucleolar maturation. Combined with the interaction of its N-terminal extension with Erb1 and the predicted RecA2-Nsa1 interaction, these findings support a model in which Drs1 coordinates early assembly events around the Nsa1-associated domains I/II with folding of domain III and ES27 around the Erb1 WD40 β-propeller at subsequent stages.

**Figure 8.**
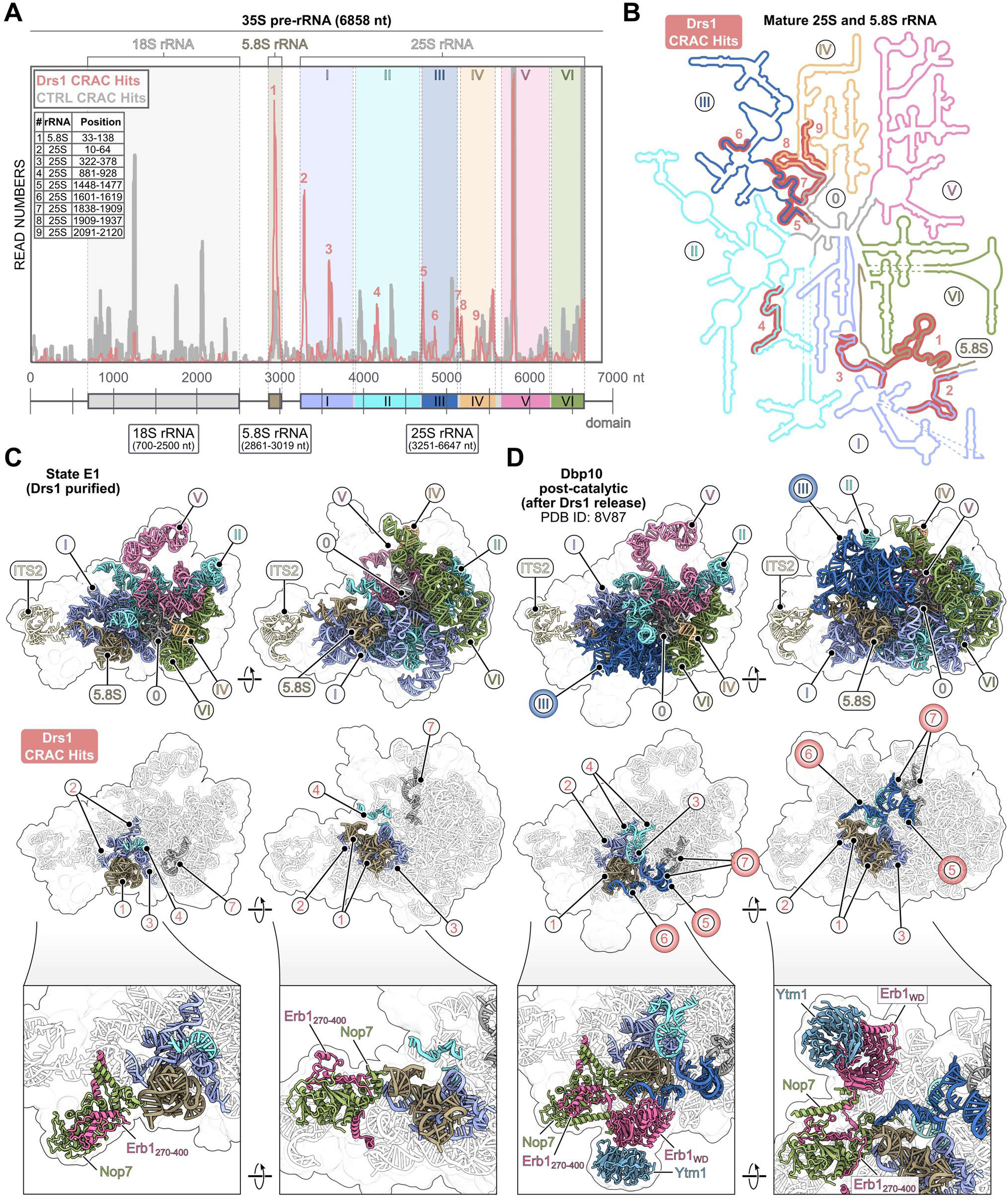
CRAC analysis of the Drs1 binding sites within pre-60S particles. (**A**) Living yeast cells expressing pATH-Drs1 or untagged control (CTRL) were UV cross-linked and processed according to the CRAC protocol. Final sequencing hits were aligned to the 35S rDNA sequence. The distribution of the pATH-Drs1 (red) and CTRL (gray) sequencing hits on the 35S rDNA is shown. Coordinates of the mature rRNA species are indicated below the graph. Cross-linking of pATH-Drs1 resulted in nine major peaks that were not detected in the CTRL experiment, exclusively localized in the 5.8S and 25S rRNA sequences, reflecting potential interaction sites of Drs1. The 5.8S rRNA and domains I–VI of the 25S rRNA are color-coded. The table shows the peak numbers (#), the corresponding rRNA species (rRNA) and the position within the rRNA species (a threshold of 5000 hits was used to define the beginning and end of each peak). (**B**) Secondary structure of the mature 25S and 5.8S rRNAs. The rRNA domains (I-VI) and the root helices (0) are labeled and color-coded as in (A). Regions of the pATH-Drs1 cross-linking hits are numbered (1-9) and highlighted in red. (**C**-**D**) Drs1 cross-linking sites within the Drs1 pre-60S State E1 (C) and within a subsequent pre-60S state after Drs1 release (D). The stably integrated rRNA domains within these states are colored and labeled as in (A-B) and shown in two orientations (upper panels). The observed crosslinking hits of Drs1 are shown below (middle panels). In the pre-60S state, after the release of Drs1 (PDB ID: 8V87), domain III of the 25S rRNA and the corresponding crosslinking hits 5-7 are highlighted (D). Close-up views highlighting the Drs1 cross-linking hits together with models for Nop7-Erb1 (C) and Ytm1-Erb1-Nop7 (D) (lower panels).

## DISCUSSION

Drs1 has long been implicated in pre-60S maturation, and in this study, we define its dual role as both a critical licensing factor for SSU-LSU separation and a remodeler of nucleolar pre-60S intermediates. Depletion or catalytic inactivation of Drs1 resulted in a pronounced accumulation of early pre-60S factors together with SSU processome components on isolated Nop7 particles. Consistently, cryo-EM analysis identified SSU processome particles almost exclusively after Drs1 depletion, providing structural support for impaired separation of emerging LSU precursors from the SSU processome. These findings place Drs1 functionally at the co-transcriptional ITS1 boundary, consistent with a role in promoting A_2_ cleavage (Talkish et al., 2016) and release of the Rrp5-Noc1-Noc2 module, which is a prerequisite for Nsa1 incorporation (Sanghai et al., 2023). Similarly, a recent study showed that depletion of another pre-60S factor, Rlp7, shifts pre-ribosomal particles toward earlier, partially co-transcriptional SSU processome states (Gerhalter et al., 2026), further highlighting the tight coupling of LSU and SSU assembly during early ribosome biogenesis. Beyond these co-transcriptional events, Drs1 is also required for productive progression of pre-60S particles through nucleolar maturation. Our cryo-EM analysis reveals a pronounced accumulation of early nucleolar intermediates upon Drs1 depletion, whereas particles containing stably incorporated 25S rRNA domain III and ES27 of domain IV and all subsequent maturation states are strongly underrepresented. Particularly prominent is an Nsa1-deficient intermediate, suggesting that loss of Drs1 function permits release of the Nsa1 module at an early stage of nucleolar maturation. Accordingly, cryo-EM structures of Drs1-associated particles span a broad series of early nucleolar intermediates but remain confined to states preceding stable domain III incorporation. Together, these complementary structural analyses define a Drs1-dependent maturation window immediately before this transition. Loss of Drs1 compromises stable association of both Nsa1 and the Nop7-Erb1-Ytm1 module, indicating that Drs1 activity promotes structural transitions that stabilize major architectural elements of the nucleolar pre-60S particle. Our CRAC data further position Drs1 at pre-rRNA regions encompassing domains III and IV, consistent with a direct role in remodeling this region. In parallel, the interaction of the Drs1 N-terminus with the C-terminal Erb1 WD40 domain links Drs1 to the Erb1-Ytm1 module, which may sense correct domain III maturation and is required, together with the helicase Spb4, for stable ES27 positioning (Cruz et al., 2024, 2022; Mitterer et al., 2023). Our biochemical reconstitution together with AlphaFold3 predictions further indicate that Drs1 and Ytm1 can simultaneously engage the Erb1 WD40 β-propeller, with the Drs1 N-terminal region positioned adjacent to the site subsequently occupied by domain III. Together, these observations suggest that Drs1 gates domain III incorporation by promoting the remodeling events required to establish a maturation-competent precursor for its stable integration. Consistent with this model, loss of Drs1 does not lead to premature domain III incorporation but instead prevents efficient progression into domain-III-containing states, thereby linking Drs1 activity to subsequent nucleolar maturation. At the spatially distant solvent side, the requirement of Drs1 for stable Nsa1 association indicates an additional role in assembly processes around domains I and II. Such coordinated activity is structurally feasible given the extended and flexible nature of the unstructured N- and C-terminal regions of Drs1, which may provide the conformational reach allowing the helicase to engage spatially separated assembly factors and pre-rRNA regions within the same pre-ribosomal particle. Together, these observations suggest that Drs1 functions as a molecular bridge, linking core pre-rRNA remodeling at the subunit interface with stabilization of solvent-exposed assembly factors. In this way, Drs1 coordinates structural transitions across distinct regions of the nucleolar pre-60S particle and promotes its orderly maturation.

Our findings place Drs1 within a broader network of DEAD-box ATPases that remodel distinct rRNA regions during LSU biogenesis. Several helicases act very early, including Dbp6, which, as part of the Npa1 complex, promotes folding of 25S rRNA root helices and also contributes to modulation of snoRNA-rRNA interactions at the immature peptidyl-transferase center (PTC) (Hamze et al., 2025; Joret et al., 2018; Khreiss et al., 2023). Dbp7 also acts at the PTC and mediates compaction of domains V and VI (Aquino et al., 2021), while Mak5, similar to Drs1, is required for efficient disconnection of primordial pre-60S particles from the SSU processome (Ismail et al., 2022) and has also been suggested to contribute to subsequent maturation steps (Brüning et al., 2018). Dbp10 facilitates maturation of the evolving PTC, where it remodels helices H89-H92, and has an additional role in restructuring domain IV H61 (Cruz et al., 2024; Manikas et al., 2016; Mitterer et al., 2024; Vanden Broeck and Klinge, 2023). Interestingly, some Dbp7 and Dbp10 binding sites in the PTC overlap, suggesting that multiple helicases act sequentially or in a partially redundant manner within the same rRNA neighborhood. A further hotspot appears to lie at the base of domain IV, where H62/H63 and ES27 undergo repeated remodeling. Spb4 catalyzes restructuring of this region during a very late nucleolar step (Brüning et al., 2018; Cruz et al., 2022; Mitterer et al., 2023), while Dbp10 remodels the neighboring root helix H61 (Cruz et al., 2024; Vanden Broeck and Klinge, 2023). Our data place Drs1 upstream of these domain IV maturation steps, at a transition characterized by stable incorporation of domain III and ES27 of domain IV. Drs1 may therefore prepare early nucleolar intermediates for subsequent remodeling events by Spb4 and, eventually, the AAA-ATPase Rea1 (Bassler et al., 2010; Cruz et al., 2022; Mitterer et al., 2023). Together, these observations highlight how successive ATP-dependent remodeling factors act on overlapping structural neighborhoods at distinct stages to enforce the ordered maturation of pre-60S particles.

## METHODS

### Yeast strains and plasmids

*Saccharomyces cerevisiae* strains used in this study are listed in **Supplementary Table 2** and are derived from the W303 background (Thomas and Rothstein, 1989). Strains were constructed using established gene disruption and genomic tagging methods (Janke et al., 2004; Longtine et al., 1998). For yeast two-hybrid analyses, the reporter strain PJ69-4A was used (James et al., 1996). Plasmids used in this study are listed in **Supplementary Table 3** and were constructed using standard DNA cloning techniques and verified by sequencing.

### Growth analyses of mutant *drs1* yeast strains

To investigate growth phenotypes upon auxin-induced depletion of Drs1, strains expressing endogenous AID-HA-*DRS1* were spotted in tenfold serial dilutions onto YPD plates in the absence or presence of auxin. Auxin-containing YPD plates were supplemented with 2 mM indole-3-acetic acid (IAA) (Sigma-Aldrich, Cat# I3750) to induce proteasomal degradation of AID-tagged Drs1, and plates were incubated at 30 °C for 2 days. To analyze growth phenotypes of plasmid-derived *drs1* mutant alleles, the AID-HA-*DRS1* strain was transformed with plasmids carrying the respective Drs1 constructs expressed under control of the endogenous *DRS1* promoter, and growth was assessed on auxin-containing YPD plates. To analyze growth phenotypes upon overexpression of Drs1 variants, strains were transformed with plasmids carrying the respective constructs under control of the galactose-inducible *GAL1-10* promoter. Transformants were spotted in tenfold serial dilutions onto SDC plates lacking leucine and containing glucose (repressing conditions) or galactose (inducing conditions) and incubated at 30 °C for the indicated times to assess growth. For plasmid-shuffle assays, a *DRS1* shuffle strain (*drs1*Δ YCplac33-P*DRS1*-*DRS1*) was transformed with plasmids carrying the indicated *drs1* mutant alleles under control of the endogenous *DRS1* promoter. Growth phenotypes were analyzed on plates containing 1 g/l 5-fluoroorotic acid (5-FOA) to select for cells that had lost the wild-type *DRS1*-containing *URA3* plasmid.

### Yeast two-hybrid analyses

Plasmids expressing the indicated Drs1 bait protein constructs from *S. cerevisiae* or *C. thermophilum*, N-terminally fused to the GAL4 DNA-binding domain (G4-BD), and Erb1 or Ytm1 prey protein constructs, N-terminally fused to the GAL4 activation domain (G4-AD), were co-transformed into the reporter strain (James et al., 1996). Yeast two-hybrid interactions were documented by spotting representative transformants in tenfold serial dilutions onto SDC-Leu-Trp (control), SDC-Leu-Trp-His (*HIS3* reporter gene), and SDC-Leu-Trp-Ade (*ADE2* reporter gene) plates and incubating at 30 °C for indicated times.

### Polysome profile analyses

Cells expressing chromosomal N-terminal fusions of Drs1 tagged with AID-HA (AID-HA-*DRS1*) were grown in YPD medium to early logarithmic growth phase. Prior to harvesting, cultures were incubated with 0.5 mM auxin for 120 min to induce proteasomal degradation of AID-HA-Drs1 or left untreated. Subsequently, 100 μg/ml cycloheximide was added and after incubation for 10 min on ice, cells were pelleted and washed once with lysis buffer [10 mM Tris-HCl (pH 7.5), 100 mM NaCl, 30 mM MgCl2, 100 μg/ml cycloheximide]. After resuspension in 3 volumes of lysis buffer and cell lysis with glass beads, the lysate was centrifuged for 10 min at 13,000 rpm at 4 °C and 6 A_260_ units of the cell extracts were loaded onto continuous 10–50% sucrose gradients (w/v) [dissolved in 50 mM Tris-HCl (pH 7.5), 100 mM NaCl, 30 mM MgCl_2_] and centrifuged with a SW40 rotor (Beckman Coulter) at 39,000 rpm for 2 h 45 min at 4 °C. Gradients were analyzed on a Foxy Jr. fraction collector (Teledyne ISCO) with continuous monitoring at 254 nm.

### Tandem affinity purification of pre-ribosomal particles

For two-step affinity purifications from yeast, respective C-terminally Flag-TEV-proteinA (FpA) or N-terminally TAP-Flag (TAPF)-tagged bait proteins were expressed under control of their native promoter. Yeast strains carrying only chromosomally encoded genes were grown in YPD, whereas strains expressing plasmid-derived bait proteins (**Figure 1G**) or plasmid-derived mutants (**Figure 3B**) were grown in synthetic dextrose complete (SDC) medium lacking leucine for plasmid selection. For depletion of auxin-inducible degron (AID) tagged chromosomal *DRS1*, cultures were incubated in the presence of 0.5 mM auxin (3-indoleacetic acid, Sigma-Aldrich, Cat# I2886) for 120 min prior to harvesting the cells. All strains were grown at 30 °C and harvested in the logarithmic growth phase, flash frozen in liquid nitrogen and stored at −20 °C. Cell pellets were resuspended in lysis buffer containing 50 mM Tris-HCl (pH 7.5), 100 mM NaCl, 5 mM MgCl_2_, 0.05% NP-40, 1 mM DTT, supplemented with 1 × SIGMAFAST protease inhibitor (Sigma-Aldrich), and cells were lysed by mechanical disruption using glass beads. Lysates were cleared by two subsequent centrifugation steps at 4 °C for 10 and 30 min at 5,000 and 16,000 rpm, respectively. Supernatants were incubated with immunoglobulin G (IgG) Sepharose 6 Fast Flow beads (GE Healthcare) on a rotating wheel at 4 °C for 90 min. Beads were transferred into Mobicol columns (MoBiTec) and, after washing with 12 ml of lysis buffer (per 2L culture volume), cleavage with tobacco etch virus (TEV) protease was performed at RT for 60 min. In a second purification step, TEV eluates were incubated with Flag agarose beads (ANTI-FlagM2 Affinity Gel, Sigma-Aldrich, Cat #A2220) for 80 min at 4 °C. After washing with 5 ml of lysis buffer, bound proteins were eluted with lysis buffer containing 300 µg/ml Flag peptide (Sigma-Aldrich, Cat# F3290) at 4 °C for 45 min. Flag eluates were analyzed by SDS-PAGE on 4-12% polyacrylamide gels (NuPAGE, Invitrogen, Cat #NP0322BOX) with colloidal Coomassie staining (Roti-blue, Carl Roth, Cat #A152.1) or by western blotting with antibodies, as indicated in the respective figures.

### Sucrose gradient centrifugation of Nsa1- and Nop7-associated particles

FpA-tagged Nsa1 and Nop7 were affinity purified as described above from cells grown in the presence or absence of 0.5 mM auxin to induce depletion of AID-HA-Drs1. The final eluates were loaded onto 10%–30% (Nsa1 eluates) or 10%–40% (Nop7 eluates) sucrose gradients (w/v) containing 50 mM Tris-HCl (pH 7.5), 100 mM NaCl, 5 mM MgCl_₂_, 0.05% NP-40 and 0.5 mM DTT, and centrifuged at 27,000 rpm for 16 h at 4 °C. Gradients were fractionated into 13 fractions and proteins were subsequently precipitated by addition of trichloroacetic acid (TCA) to a final concentration of 15%. TCA-precipitated proteins were resuspended in SDS sample buffer and analyzed by SDS-PAGE on 4-12% polyacrylamide gels followed by Coomassie blue staining or western blotting.

### Western blotting

Western blot analysis was performed using the following antibodies: anti-Nog1 antibody (1:5,000), anti-Nog2 antibody (1:5,000), anti-Nsa2 antibody (1:5,000), provided by Micheline Fromont-Racine; anti-Nug1 antibody (1:10,000), anti-Bud20 antibody (1:5,000), anti-Yvh1/anti-Nop7 antibody (1:4,000) provided by Vikram Panse; anti-Has1 antibody (1:10,000) provided by Patrick Linder; anti-Ebp2 antibody (1:10,000), provided by Keiko Mizuta; anti-Rpl3 antibody (1:5,000), provided by Jonathan Warner; anti-Rps3 antibody (1:30,000) provided by Matthias Seedorf; anti-GAPDH antibody (1:30,000; Thermo Fisher Scientific, Cat# MA5-17538 RRID:AB_10977387), horseradish-peroxidase-conjugated anti-Flag antibody (1:10,000; Sigma-Aldrich, Cat# A8592, RRID:AB_439702), horseradish-peroxidase-conjugated anti-HA antibody (1:5,000; Roche, Cat# 12013819001; RRID:AB_390917), secondary horseradish-peroxidase-conjugated goat anti-rabbit antibody (1:2,000; Bio-Rad, Cat# 166-2408EDU, RRID:AB_11125345), secondary horseradish-peroxidase-conjugated goat anti-mouse antibody (1:2,000; Bio-Rad, Cat# STAR105P, RRID:AB_323002).

### Purification of Nop7-FpA particles for cryo-EM analysis

*NOP7*-FpA AID-HA-*DRS1* cells were grown in YPD medium. Depletion of chromosomally tagged Aid-HA-Drs1 was induced by adding 500 µM auxin (final concentration, Sigma-Aldrich) for 90 minutes. The cells were harvested at an OD_600_ of approximately 2.5 and flash frozen at −80 °C. Lysis was carried out using a SPEX 6970EFM Freezer/Mill. The cell powder was resuspended in buffer (60 mM Tris, pH 7.5, 50 mM NaCl, 40 mM KCl, 5 mM MgCl₂, 1 mM DTT) supplemented with 5% glycerol, 0.1% NP-40 and Complete EDTA-free protease inhibitor (Roche). The lysate was cleared by centrifugation, first for 15 min at 4,000 rpm (Eppendorf 5810R Centrifuge) and then for 25 min at 17,500 rpm (Sorvall LYNX 6000 Superspeed Centrifuge), both at 4 °C. Magnetic Dynabeads M-270 (Invitrogen), which were coated with IgG, were then added to the cleared lysate and incubated under rotation for 90 minutes at 4 °C. The beads were then washed, first with buffer supplemented with 2% glycerol and 0.01% NP-40, and second with buffer supplemented with 2% glycerol and 0.05% β-octylglucoside. Samples were eluted by the addition of homemade TEV protease and incubation for 90 minutes at 16 °C. The samples were centrifuged for 5 minutes at 14,000 rpm in a tabletop centrifuge at 4 °C, and the supernatant was used for cryo-EM grid preparation.

### Purification of Drs1 particles for cryo-EM analysis

The AID-HA-*DRS1* strain was transformed with the respective YCplac111-P.*DRS1*-Drs1 plasmids (wt, K284A, E389Q). Cells were grown overnight in SDC-Leu media and the next day shifted to YPD media. Cells were grown for about 4.5 hours and depletion of the endogenous tagged AID-HA-Drs1 copy was induced by the addition of 500 µM Auxin (final concentration, Sigma-Aldrich) for 90 min. Cells were harvested by centrifugation, flash frozen in liquid nitrogen and stored at −80

°C. Cells were disrupted with a SPEX 6970EFM Freezer/Mill. The cell powder was resuspended in buffer (50 mM Tris-HCl, pH 7.5, 100 mM NaCl, 5 mM MgCl_2_, 1 mM DTT) supplemented with 5% glycerol, 0.1% NP-40 and Complete EDTA-free protease inhibitor (Roche). The lysate was cleared twice by centrifugation (1st at 4,000 rpm for 15 min and 2nd at 17,500 rpm for 25 min). IgG Sepharose™ 6 Fast Flow affinity resin was added to the cleared lysate and incubated for 90 minutes under rotation at 4 °C. Beads were washed with buffer supplemented with 5% glycerol and 0.1% NP-40 and the samples were eluted by incubation with homemade TEV protease for 90 min at 16 °C. The eluate was incubated with Anti-FLAG M2 agarose beads (Sigma-Aldrich) for 90 min at 4 °C and the beads were washed first with buffer supplemented with 0.01% NP-40 and then with buffer supplemented with 0.05% C12E8 (Nikkol) using 1 ml Mobicol columns (MoBiTec). Samples were eluted by the addition of 3x Flag peptide (Sigma-Aldrich) at a final concentration of 250 µg/ml. After incubation for 60 min at 4 °C, the samples were eluted by centrifugation at 2,000 rpm in a tabletop centrifuge at 4 °C. Potential aggregates were removed by centrifugation at 14,000 rpm for 5 min and 4 °C and the supernatant was used for grid preparation.

### Cryo-EM grid preparation

For all samples, 3.5 µL were applied to R3/3 300 mesh copper grids with carbon support coated with 3 nm continuous carbon (Quantifoil). Grids were blotted using a Vitrobot Mark IV (FEI Company) at 4 °C and 85-90% humidity for 3 s and with 45 s pre-blot time before plunge-freezing in liquid ethane.

### Cryo-EM data collection

The datasets of the Nop7-FpA samples with and without auxin depletion of Drs1 were collected on a Titan Krios G1 at 300 keV (Thermo Fischer) and equipped with a K2 summit direct electron detector (Gatan) with a nominal pixel size of 1.060 Å/pixel and a total dose of 45.12 e^-^/Å^2^ and 44.88 e^-^/Å^2^, respectively. EPU (Thermo Fisher) was used to collect 15,285 micrographs of the Nop7-FpA - Auxin and 12,372 micrographs of the Nop7-FpA + Auxin sample. For the TAPF-Drs1 wildtype and the Drs1 mutant samples, data was collected on a Titan Krios G3 at 300 keV (Thermo Fisher) equipped with a Falcon 4i direct electron detector (Thermo Fisher) and a Selectris X energy filter (Thermo Fisher). Data was collected with a nominal pixel size of 0.727 Å/pixel, a filter slid width of 20 eV and a total dose of 40 e^-^/Å^2^. For the TAPF-Drs1-wt, -K284A and -E389Q datasets 14,973, 16,898 and 15,600 micrographs were collected, respectively using EPU (Thermo Fisher). Data was collected with a defocus range of –0.5 to –3.5 µm. Motion correction was performed with MotionCor2 (Zheng et al., 2017) and CTFFIND4 (Rohou and Grigorieff, 2015) was used for estimation of contrast transfer function (CTF) parameters.

### Cryo-EM data image processing

For the Nop7 datasets with (-Auxin) and without (+Auxin) Drs1, 15,182 and 12,098 micrographs were selected, respectively, for cryo-EM data processing. To allow for better comparison, the datasets were combined and processed together. Particles were picked using RELION 4.0 Autopick (Kimanius et al., 2021) with the Laplacian-of-Gaussian blob detection, and 2,532,399 initial particles were extracted with a pixel size of 3.18 Å/pixel and a box size of 150 × 150 pixel. Three consecutive rounds of 2D classification were performed in cryoSPARC (Punjani et al., 2017), and the pre-60S (860,063 particles) and SSU processome (29,072 particles) class averages were selected. Following Ab-initio reconstruction and homogeneous refinement, both particle classes were imported into RELION 4.0. Reextracted particles (3.18 Å/pixel, box size 150 × 150 pixel) were then used for 3D refinement, followed by elaborate 3D classifications without alignment (see **Supplementary Figure 3** for details). The final classes were 3D refined, and the number of particles in each dataset was calculated using the unique optical group identifier.

For Drs1 wt, Drs1 K284A and Drs1 E389Q, 13,307, 15,010 and 13,500 micrographs were used, respectively. Particles were picked using RELION 4.0 Autopick using Laplacian-of-Gaussian blob detection and extracted with a pixel size of 2.181 Å/pixel (box size 200 × 200 pixel). Three rounds of 2D classification were performed in cryoSPARC for each dataset. Pre-60S 2D class averages were selected and used for ab-initio reconstructions and homogeneous refinements. 3D refinement followed by 3D classification (without alignment) was performed in RELION 4.0. All three datasets showed similar classes, ranging from early nucleolar State A/B particles (Kater et al., 2017) to more evolved nuclear pre-60S particles, which contain stable uL1 stalk binding, Nip7-Nop2 and RNA helicase Dbp10, for example. However, all particle classes lacked the 25S rRNA domain III and were pre-60S precursors prior to State D (Kater et al., 2017). The Drs1 K284A and Drs1 E389Q mutant datasets also contained particle classes lacking the Nsa1 module (Nsa1, Rpf1, Mak16 and Rrp1). No additional density corresponding to Drs1 could be observed for any of the datasets or particle classes. To increase the number of particles for the individual pre-60S states and enable better comparison of the datasets, we combined them for unbiased data processing. As with the individual datasets, particles were picked using RELION 4.0 Autopick. This yielded a total of 2,042,641 particles (Drs1 WT: 670,231; Drs1 K284A: 798,734; Drs1 E389Q: 573,676), which were extracted with a pixel size of 2.181 Å/pixel (box size 200 × 200 pixel). These were used for three rounds of 2D classification in cryoSPARC. Selected 2D class averages were then used for Ab-initio reconstruction and homogeneous refinement, after which the particles were reimported into RELION 4.0. All subsequent 3D classifications and focused 3D classifications were performed in RELION 4.0, without alignment. After an initial 3D classification step with ten classes, three classes containing poor-quality particles were discarded; the remaining seven classes were arranged according to their maturation status and used for extensive 3D classifications. The particle numbers and percentage of the individual datasets were calculated for all obtained classes (see **Supplementary Figure 5-6** for details). Classes used for model building were extracted with a pixel size of 0.727 Å/pixel and a box size of 640 × 640 pixel and reimported into cryoSPARC for non-uniform refinement, including defocus and global CTF refinement. Masks were used for particle subtraction and focused local refinements of flexible modules within the different states. The local refinements were fitted into the consensus map and composite maps were generated with ChimeraX using the vop max command (Pettersen et al., 2021).

### Model building

The molecular models of nucleolar pre-60S states, including State B (PDB ID: 6EM4) (Kater et al., 2017), State 2 (PDB ID: 6C0F) (Sanghai et al., 2018) and Dbp10 pre-catalytic State (PDB ID: 8V84) (Cruz et al., 2024) were used as initial models for the best resolved Drs1 pre-60S State B2 and were fitted into the cryo-EM density map with ChimeraX (Pettersen et al., 2021). Model building was assisted by AF2 and AF3 predictions (Abramson et al., 2024; Jumper et al., 2021). All model building and adjustments were performed using Coot (v 0.9.8.96) (Emsley et al., 2010). The State B2 model was used as starting model for States A and B1 and fitted into the densities. Parts of the model missing in the respective states were removed, and the models were manually adjusted in Coot. For State E1, initial models included the Drs1 State B2, Dbp10 post-catalytic (PDB ID: 8V87) (Cruz et al., 2024), Dbp10-2 (PDB ID: 8I9V) (Cruz et al., 2024), State E2 (PDB ID: 7NAC) (Cruz et al., 2022) and State D (PDB ID: 8BVU) (Mitterer et al., 2023). Drs1 States E1 and B2 were used to generate the models of States C, E2, and D1/D2. For State C, the molecular model of the PTC helices H89-92 from the Dbp10 pre-catalytic State (PDB ID: 8V84) (Cruz et al., 2024) was fitted into the density and adjusted in Coot. Molecular models were real space refined with Phenix (v 1.19.2) using secondary structure restraints (Liebschner et al., 2019) and validated with MolProbity (Chen et al., 2010).

Figures of cryo-EM density maps and molecular models were visualized with ChimeraX (Pettersen et al., 2021).

### Co-expression and interaction studies of *Ct*Drs1 and *Ct*Erb1 constructs

Yeast cells were transformed with two *GAL1-10* overexpression plasmids with *TRP1* and *LEU2* selection markers, respectively (see **Supplementary Table 3**). Cells were grown in SRC-Leu-Trp media until they reached an OD600 of 2.5-3.0. Expression was induced by the addition of an equal volume of 2xYPG (4% galactose) media, and cells were grown for 4 h and harvested by centrifugation. Cells were lysed with a SPEX 6970EFM Freezer/Mill and the cell powder was resuspended in buffer containing 20 mM HEPES, pH 7.5, 150 mM NaCl, 10 mM _MgCl2_, 10 mM KCl, 5% glycerol, 0.1% NP-40 and 1 mM DTT. Lysates were cleared by centrifugation and incubated for 90 min with IgG Sepharose™ 6 Fast Flow affinity resin. The IgG beads were washed, and samples were eluted with TEV protease. Anti-FLAG M2 agarose beads (Sigma-Aldrich) were added to the eluate and incubated for 90 min at 4 °C on a rotating wheel. Beads were washed with 10 mL of buffer using 1 ml Mobicol columns (MoBiTec) column and eluted with buffer supplemented with 3x Flag peptide (final concentration 250 µg/ml, Sigma-Aldrich). Samples were analyzed by SDS-PAGE and Coomassie staining, as well as Western blotting.

Expression and purification of *C. thermophilum* proteins

The *Ct*Ytm1-*Ct*Erb1_WD_ complex was purified as previously described (Thoms et al., 2016), with an additional gel filtration step in binding buffer (20 mM HEPES-KOH pH 7.5; 200 mM NaCl; 10 mM MgCl_₂_; 10 mM KCl; 1 mM DTT; 5% glycerol) at the end of the purification. *Ct*Drs1_1-129_ was overexpressed in *E. coli* Rosetta 2 (DE3) cells by adding 0.25 mM IPTG at an OD_600_ of 0.8–1.0, followed by an 18-hour incubation at 18 °C. The cells were then harvested and resuspended in buffer A (20 mM Tris-HCl pH 7.5, 500 mM NaCl and 1 mM DTT), supplemented with a protease inhibitor cocktail. The resuspended cells were lysed by passing them through a cell disruptor (Constant Systems Ltd.) once at 2000 bar. The supernatant was obtained by centrifugation and supplemented with 10 mM imidazole. The first affinity purification was conducted using Ni-NTA agarose beads (Macherey-Nagel). The eluate was then desalted using buffer B (50 mM Tris-HCl pH 7.5, 150 mM NaCl and 1 mM DTT). The poly-histidine tag was then cleaved with TEV protease and removed by passing the solution through a Ni-NTA column. The proteins were further purified by anion exchange chromatography using Q Sepharose™ Fast Flow resin (Cytiva). Elution was performed in steps, gradually increasing the NaCl concentration of the buffer (20 mM HEPES-KOH pH 7.5, 100 mM NaCl, 10 mM MgCl₂, 10 mM KCl, 1 mM DTT, 5% glycerol) up to 1 M. Finally, the fractions containing the protein were pooled and applied to a Superdex 200 Increase 10/300 GL column (Cytiva). The peak fractions were then pooled, concentrated, and flash-frozen before being stored at −80 °C until use.

### Binding assay of the *Ct*Ytm1-*Ct*Erb1_WD_ complex with *Ct*Drs1_1-129_

*Ct*Ytm1-*Ct*Erb1_WD_ (550 pmol) and *Ct*Drs1_1-129_ (1,100 pmol) were mixed in 50 µl of binding buffer (20 mM HEPES-KOH pH 7.5, 200 mM NaCl, 10 mM MgCl_₂_, 10 mM KCl, 1 mM DTT, 5% glycerol) and incubated on ice for 10 minutes. The sample was then injected into a Superose 6 Increase 3.2/300 column equilibrated with binding buffer. Peak fractions were analyzed by SDS-PAGE and Coomassie blue staining. Samples containing *Ct*Ytm1-*Ct*Erb1_WD_ and *Ct*Drs1_1-129_ alone were also prepared and analyzed in the same way.

### CRAC protocol

Yeast cells expressing Drs1 fused at the N-terminus to proteinA-TEV-(His)_6_ (pATH) and the wild-type BY4742 strain (negative control) were grown at 30 °C in 2.8 L of SDC-Trp medium to an OD_600_ of 0.6. Cells were irradiated for 100 s with UV light at 254 nm and harvested. Cells were resuspended in TNM150 buffer (50 mM Tris-HCl, pH 7.8, 150 mM NaCl, 1.5 mM MgCl_2_, 0.1% NP-40, 5 mM β-mercaptoethanol) and lysed by mechanical disruption using zirconia beads. Cell lysates were mixed with 400 μL IgG Sepharose™ 6 Fast Flow slurry (GE Healthcare) pre-equilibrated with TNM150 buffer and incubated for two hours at 4 °C on a stirring wheel. Beads were washed two times with TNM1000 buffer (50 mM Tris–HCl (pH 7.8), 1 M NaCl, 1.5 mM MgCl_2_, 0.1% NP-40, 5 mM β-mercaptoethanol) and two times with TNM150 buffer, then resuspended in 600 μL TNM150 buffer and transferred into Micro Bio-Spin 6 columns (Bio-Rad). Pre-ribosome elution from IgG Sepharose beads was achieved by incubation with 30 μL homemade GST-tagged TEV protease for two hours on a shaking table at 16 °C. TEV eluates (about 650– 700 μL) were partially digested for 5 min at 37 °C with 1.4 μL of RNace-IT (Agilent) diluted to 1:50 in TNM150 buffer and the reactions were stopped using 0.4 g of guanidine hydrochloride. The resulting samples were supplemented with 300 mM NaCl and 10 mM imidazole and incubated overnight at 4 °C on a stirring wheel with 50 μl Ni-NTA agarose resin slurry (QIAGEN) pre-equilibrated with wash buffer I (50 mM Tris–HCl, pH 7.8, 300 mM NaCl, 10 mM imidazole, 6 M guanidine hydrochloride, 0.1% NP-40, 5 mM β-mercaptoethanol). Ni-NTA beads were then washed two times with wash buffer I, three times with 1× PNK buffer (50 mM Tris–HCl, pH 7.8, 10 mM MgCl_2_, 0.5% NP-40, 5 mM β-mercaptoethanol) and transferred into Pierce Spin Columns (Thermo Scientific). RNAs retained on the Ni-NTA beads were dephosphorylated for 30 min at 37 °C using TSAP (Promega) in 1× PNK buffer in a total volume of 80 μL in the presence of 80 units of RNasin ribonuclease inhibitor (Promega). Beads were washed once with wash buffer I to inactivate TSAP and three times with 1× PNK buffer. The miRCat-33 linker (5′-AppTGG AAT TCT CGG GTG CCA AG/ddC/-3′) was ligated to the 3′ end of the RNAs on the Ni-NTA beads with 800 units of T4 RNA ligase 2 truncated K227Q (New England Biolabs) in 1 x PNK buffer / 16.67% PEG 8000 in the presence of 80 units RNasin in a total volume of 80 μL. The ligation reaction was incubated for five hours at 25 °C. Beads were washed once with wash buffer I to inactivate the RNA ligase and 3 times with 1× PNK buffer. The 5′ ends of the RNAs were then radiolabeled by phosphorylation in reactions containing 1× PNK buffer, 40 μCi of 32P-ɣ ATP and 20 units of T4 PNK (Sigma) in a total volume of 80 μL. The reactions were incubated at 37 °C for 40 min. To ensure all RNAs get phosphorylated at the 5′ end for downstream ligation of the 5′ linker, 1 μL of 100 mM ATP was added to the reaction mix, which was incubated for another 20 min at 37 °C. Beads were washed once with wash buffer I to inactivate the kinase and four times with 1 x PNK buffer. Solexa linkers L5Aa (5′-invddT-ACA CrGrAr CrGrCr UrCrUr UrCrCr GrArUr CrUrNr NrNrUr ArArG rC-OH-3′) and L5Ac (5′-invddT-ACA CrGrAr CrGrCr UrCrUr UrCrCr GrArUr CrUrNr NrNrGr CrGrCr ArGrC-OH-3′) were ligated to the 5′ end of the RNAs retained on the Ni-NTA beads for the Drs1-HTpA and BY4742 samples, respectively. Ligation reactions contained 1× PNK buffer, 1.25 μM of the relevant Solexa linker, 1 mM ATP, 40 units of T4 RNA Ligase 1 (New England Biolabs) in a total volume of 80 μL. The reactions were incubated overnight at 16 °C. Beads were washed three times with wash buffer II (50 mM Tris–HCl, pH 7.8, 50 mM NaCl, 10 mM imidazole, 0.1% NP-40, 5 mM β-mercaptoethanol). The material retained on the beads was then eluted using two times 200 μL elution buffer (50 mM Tris-HCl, pH 7.8, 50 mM NaCl, 150 mM imidazole, 0.1% NP-40, 5 mM β-mercaptoethanol). Eluates were precipitated with TCA (20% final concentration) in the presence of 30 μg of glycogen (Roche) to favor precipitation. The precipitated material was resuspended in NuPAGE™ LDS sample buffer (Invitrogen) with reducing agent (Invitrogen), heated 10 min at 65 °C, loaded on NuPAGE™ 4–12% Bis–Tris gels (Invitrogen) and run in 1× MOPS SDS running buffer (Invitrogen). The material was then transferred onto Amersham Protran Nitrocellulose Blotting Membrane (GE Healthcare), using a transfer buffer containing 1× NuPAGE (Invitrogen) and 20% MeOH, for two hours at 25 V and 4 °C. For the pATH-Drs1 sample, the area of the membrane containing a radioactive signal at the expected size of His_(6)_-Drs1 protein was excised. A membrane area at the same size was excised in the BY4742 sample lane. Membranes were soaked in 400 μl wash buffer II supplemented with 1% SDS, 5 mM EDTA and proteins were degraded using 100 μg proteinase K (Sigma) and incubation for two hours at 55 °C. RNA was extracted with phenol:chloroform:isoamyl alcohol (25:24:1) and then precipitated by addition of 1:10 volume of 3 M sodium acetate (pH 5.2), 2.5 volumes of 100% ethanol and 20 μg of glycogen. Dried RNA pellets were dissolved in ultrapure MilliQ H2O. Synthesis of cDNAs was performed using SuperScript III reverse transcriptase (Thermo Fisher Scientific) and oligonucleotide miRcatRT (5′-CCT TGG CAC CCG AGA ATT-3′). The resulting cDNAs were PCR-amplified using LA Taq DNA polymerase (TaKaRa) and primers P5F (5′-AAT GAT ACG GCG ACC ACC GAG ATC TAC ACT CTT TCC CTA CAC GAC GCT CTT CCG ATC T-3′) and P3R (5′-CAA GCA GAA GAC GGC ATA CGA GAT CCT TGG CAC CCG AGA ATT CC-3′). The resulting PCR products were purified by phenol:chloroform:isoamyl alcohol extraction and ethanol precipitation. After agarose gel electrophoresis (agarose ‘small fragments’, Eurogentec) run in 1× TBE buffer and stained with SYBR Safe DNA gel stain (Invitrogen), DNA fragments ranging in size between 150 and 250 base pairs were gel purified using MinElute PCR Purification Kit (QIAGEN). Concentration of the final DNA samples was measured using Qubit™ dsDNA HS Assay Kit (Invitrogen) and a Qubit™ fluorometer (Thermo Fisher Scientific) and the samples were sent to the GeT-PlaGe Genotoul facility for Illumina sequencing.

## DATA AVAILABILITY STATEMENT

The data underlying this study is available in the article and in its online supplementary material. The final cryo-EM composite maps and molecular models were deposited to the Electron Microscopy Data Bank (EMDB) and the Protein Data Bank (PDB) with the accession codes: EMD-XXXXX and XXXX for State A; EMD-XXXXX and XXXX for State B1; EMD-XXXXX and XXXX for State B2; EMD-XXXXX and XXXX for State C; EMD-XXXXX and XXXX for State D1; EMD-XXXXX and XXXX for State D2; EMD-XXXXX and XXXX for State E1; EMD-XXXXX and XXXX for State E2. The consensus maps, as well as local refinement maps used to generate composite maps, were also deposited to the EMDB and are listed in **Supplementary Table 1**. The NGS analysis data (CRAC experiment) were deposited to the Gene Expression Omnibus database with the accession number GSEXXXXXX.

## FUNDING

This research was funded in whole or in part by the Austrian Science Fund (FWF) grants [10.55776/P37114] and [10.55776/PAT8544624] to V.M., the ERC grant [ADG 741781 GLOWSOME] to E.H., and the ERC grants [ADG 885711 HumanRibogenesis] and [SYG 101224119 snoOPERA] to R.B., the ANR grants “RIBOPRE60S” and “DUKKED” to A.K.H., and the ANR grant ANR-21-CE12-0008-01 to B.A. The thesis of H.H. was funded by the French Ministry of Higher Education and Research and the “Ligue Nationale Contre le Cancer”. T.D. was supported by the Graduate School of Quantitative Biosciences Munich (QBM). RB was funded by DFG - Project-ID 533767322 - EXC 3113/1, Cluster for Nucleic Acid Sciences and Technologies - NUCLEATE. For open access purposes, the authors have applied a CC BY public copyright license to any author accepted manuscript version arising from this submission.

### AUTHOR CONTRIBUTIONS

Conceptualization: M.T., V.M. Resources: M.T., S.A.K., K.A., A.T., V.M. Investigation: M.T., S.A.K., K.A., H.H., A.T., N.M.V., A.K.H., V.M. Formal analysis & Validation: M.T., S.A.K., K.A., H.H., A.T., T.D., B.A., L.C., A.K.H., R.B., V.M. Visualization: M.T., S.A.K., K.A., A.T., T.D., N.M.V., L.C., A.K.H., V.M.; Supervision: M.T., E.H., A.K.H., R.B., V.M. Funding acquisition: E.H., A.K.H., R.B., V.M. Writing – original draft preparation: V.M. Writing – review & editing: M.T., S.A.K., K.A., H.H., A.T., T.D., B.A., N.M.V., L.C., E.L., E.H., A.K.H., R.B., V.M.

## ACKNOWLEDGMENTS

We thank Otto Berninghausen, Susanne Rieder and Charlotte Ungewickell for assistance with cryo-EM data acquisition. We thank Petra Ihrig and Jürgen Reichert from the BZH Heidelberg MS facility for performing MALDI-TOF mass spectrometry analyses. We are grateful to Virginie Marchand and Yuri Motorin (EpiRNA-Seq, IBSLor, Nancy, France) for the sequencing of CRAC libraries.

