## Supplementary Information for "RNA helicase Drs1 gates 25S rRNA domain III incorporation during early nucleolar pre-60S maturation"

6 Cluster for Nucleic Acid Sciences and Technologies - NUCLEATE

### Contributed equally

* Corresponding authors

**Supplementary Information associated with this article includes:**

Supplementary Figure 1-12

Supplementary Table 1-3

Supplementary References

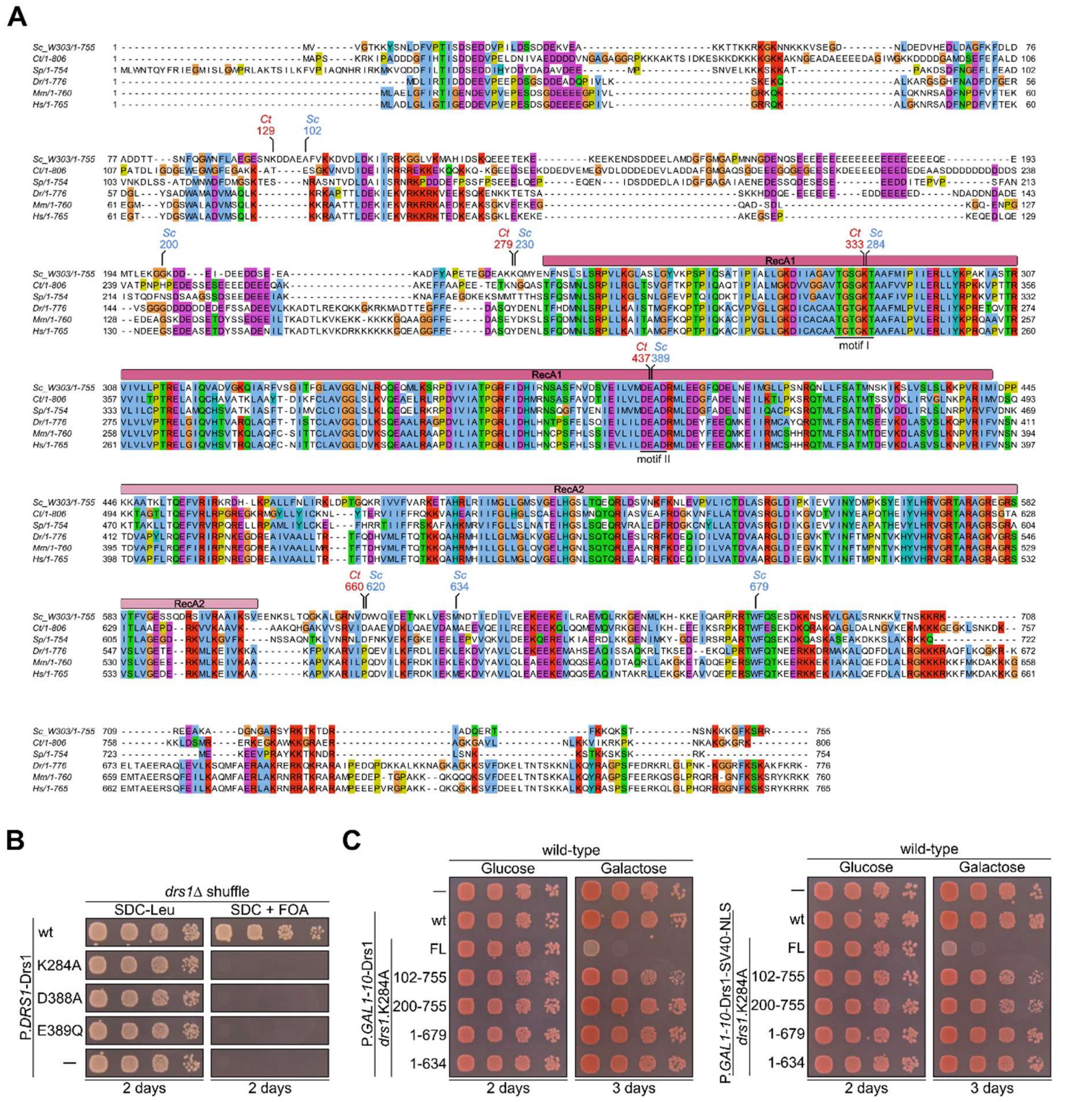

**Supplementary Fig. 1. The Drs1 catalytic core and terminal extensions are required for Drs1 function.** (**A**) The amino acid sequences of Drs1 from *Saccharomyces cerevisiae* (*Sc*, W303), *Chaetomium thermophilum* (*Ct*), *Schizosaccharomyces pombe* (*Sp*), *Danio rerio* (*Dr*), *Mus musculus* (*Mm*), and *Homo sapiens* (*Hs*) were aligned and displayed using Clustal Omega and Jalview (Madeira et al., 2024; Waterhouse et al., 2009). The RecA1 and RecA2 domains and Walker A (motif I) and Walker B (motif II) motifs are indicated. Truncations and residues mutated in this study are labeled (*S. cerevisiae, Sc*; *C. thermophilum, Ct*). (**B**) The *drs1*Δ shuffle strain was transformed with plasmids harboring wild-type (wt) *DRS1* or indicated catalytic mutants under control of the *DRS1* promoter (empty plasmid, –). Transformants were spotted in tenfold serial dilutions on SDC-Leu (plasmid control) or SDC + FOA plates and growth was monitored after incubation for 2 days at 30 °C. (**C**) Effect of terminal Drs1 truncations on dominant-negative growth phenotype. Wild-type *DRS1*, full-length *drs1* K284A, or indicated K284A N- and C-terminal truncation mutants were overexpressed without (left panel) or with (right panel) an SV40 nuclear localization signal (NLS). Transformants were spotted in tenfold serial dilutions on glucose or galactose plates and incubated at 30 °C for the indicated times.

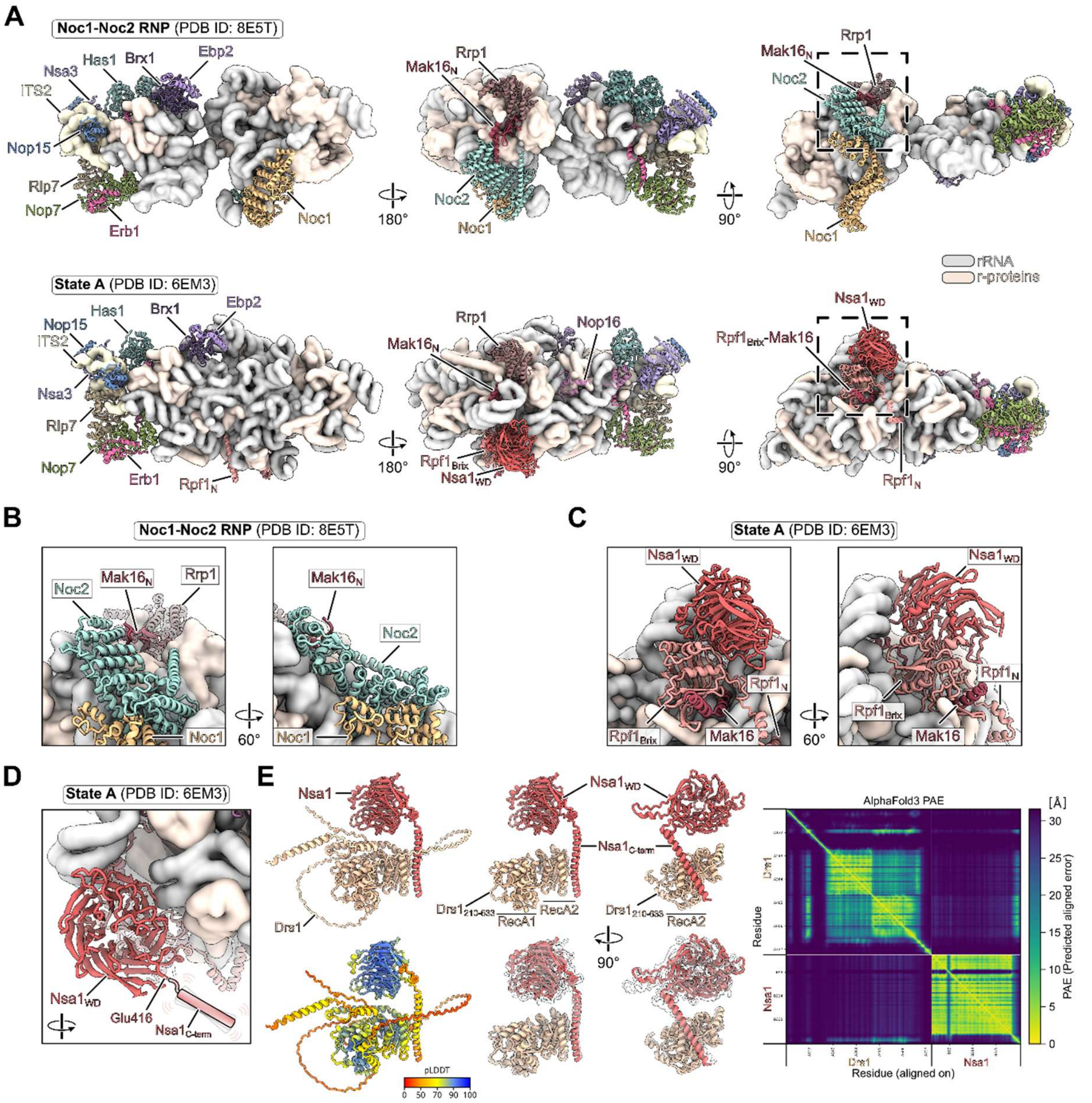

**Supplementary Fig. 2. Noc1-Noc2 blocks the Nsa1-Rpf1-Mak16 binding site on early pre-60S.** (**A**) Comparison of the early Noc1-Noc2 state (PDB ID: 8E5T, upper panels) with the subsequent State A (PDB ID: 6EM3, lower panels) shown in three orientations (Kater et al., 2017; Sanghai et al., 2023). The 25S rRNA model in State A was rigid-body fitted to the 25S rRNA domain I in the Noc1-Noc2 particle. Assembly factors are highlighted and labelled. Ribosomal proteins and rRNA are shown as filtered surface views for clarity. (**B**-**C**) Close-up views of the overlapping Noc1-Noc2 (PDB ID: 8E5T) (B) and Nsa1-Rpf1-Mak16 (PDB ID: 6EM3) binding regions (C). (**D**) Close-up view of the Nsa1 WD40 β-propeller within State A (PDB ID: 6EM3). The unresolved C-terminal α-helix is indicated (PDB ID: 6EM3). (**E**) AlphaFold3 prediction (Abramson et al., 2024) of the potential Drs1-Nsa1 interaction. The Drs1-Nsa1 complex is coloured by protein and according to the pLDDT score (left panels). The $Drs1_{210\text{-}633}$-Nsa1 complex, omitting the N- and C-terminal extensions of Drs1, is shown for clarity and overlaid with lower-ranked AF3 models (middle panels). The predicted aligned error (PAE) plot is shown (right panel).

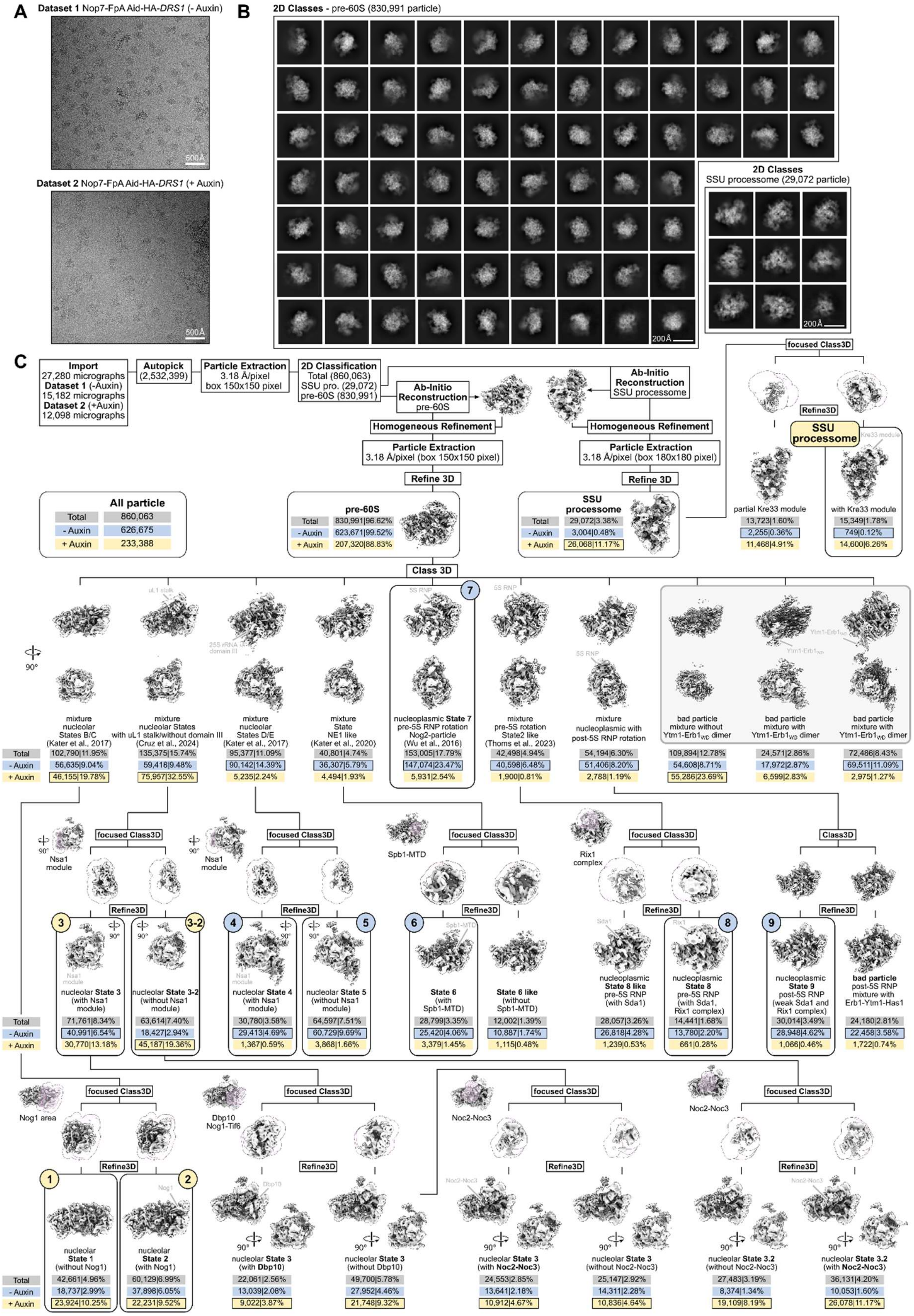

**Supplementary Fig. 3. Cryo-EM sorting scheme of the combined Nop7 datasets with and without Drs1.** (**A**) Representative electron micrographs of the Nop7-FpA Aid-HA-*DRS1* (dataset 1) and the Nop7-FpA Aid-HA-*DRS1* + Auxin (dataset 2) datasets. (**B**) 2D class averages of pre-60S and SSU processome particles after combining dataset 1 and 2. (**C**) Processing scheme of the combined dataset. The particle numbers and percentages are shown for each class as total (combined dataset) and for the parental datasets (- Auxin, with Drs1/+ Auxin, without Drs1). The pre-60S states 1-9 and the SSU processome used in **Figure 4** are highlighted. States 3 and 3-2 can be further sorted for the presence or absence of either Dbp10 or the Noc2-Noc3 dimer (lower panels).

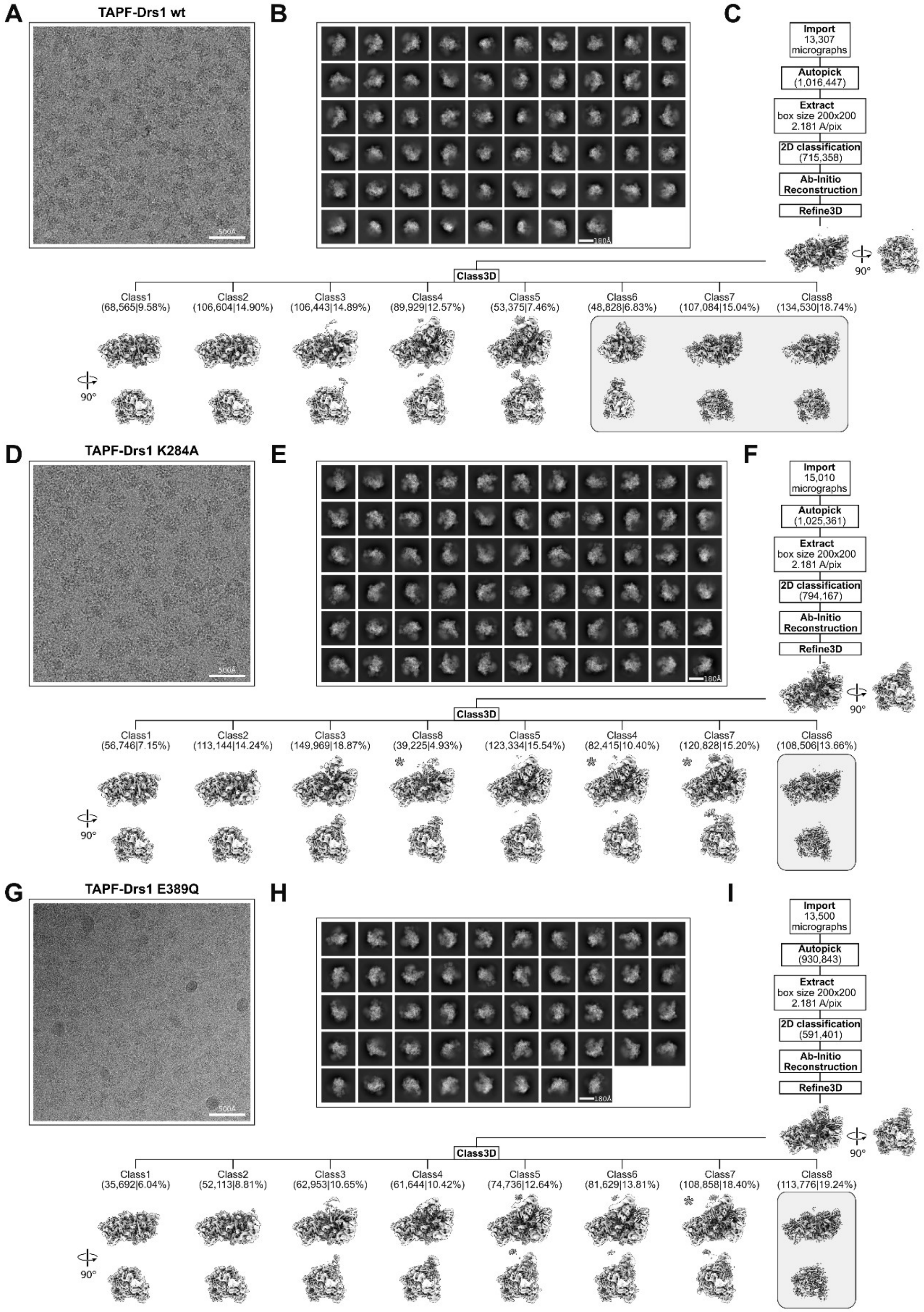

**Supplementary Fig. 4. Cryo-EM processing schemes of the individual Drs1 wt and mutant datasets.** (**A**, **D**, **G**) Representative micrographs, (**B**, **E**, **H**) 2D class averages and (**C**, **F**, **I**) processing schemes of the individual datasets. Particle numbers and percentages of the classes are shown in brackets. The particle classes are arranged from left to right according to their maturation state, from early to middle nucleolar pre-60S. Classes missing the Nsa1-module are labeled with an asterisk. Bad particle classes are indicated with gray boxes.

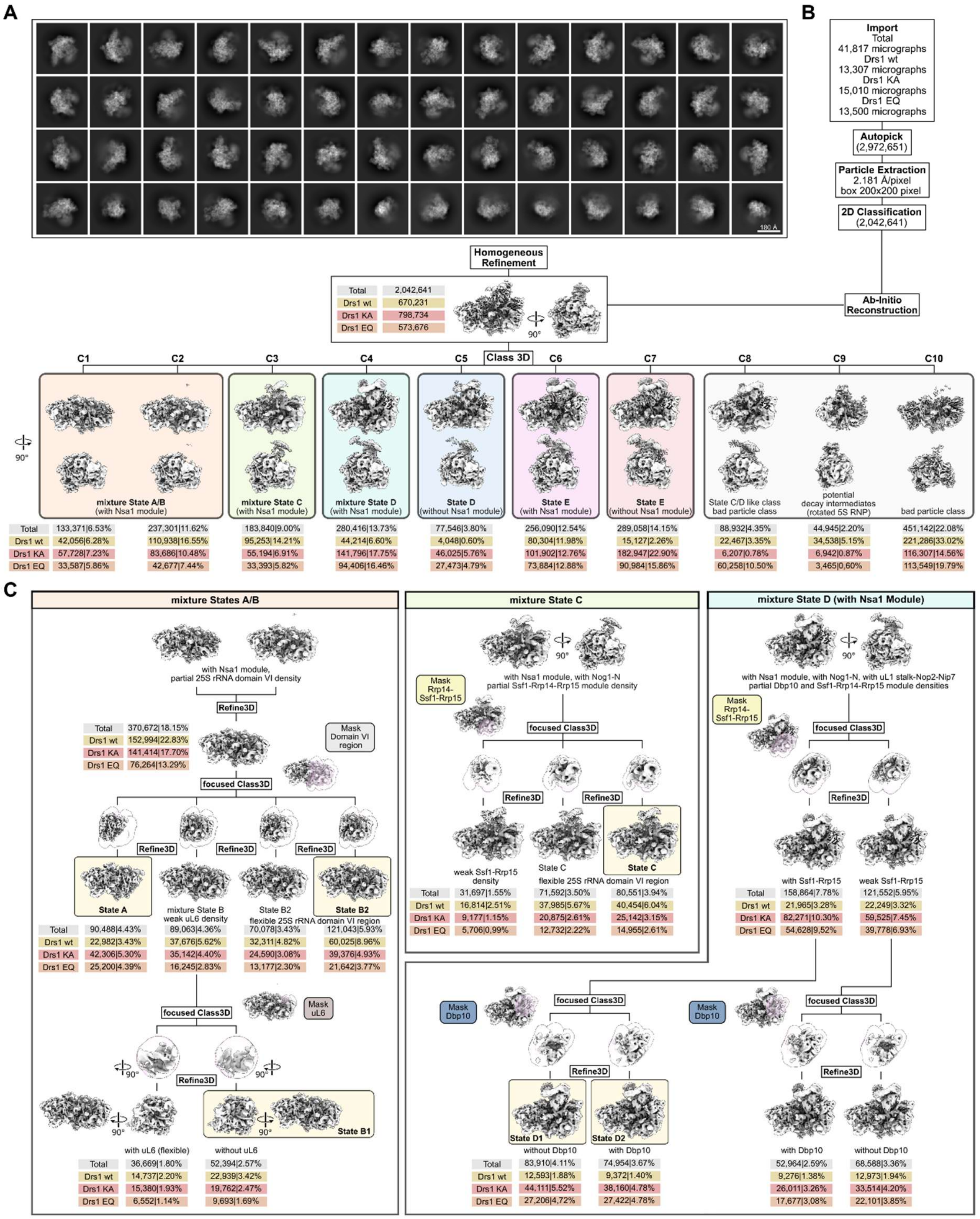

**Supplementary Fig. 5. Processing scheme of the combined Drs1 wt and mutant datasets – Part I.** (**A**) 2D class averages and (**B**) initial processing and 3D classification of the combined dataset. (**C**) Sorting schemes of the individual classes after the initial round of 3D classification (B). Particle numbers and their percentages are shown for each class as total (combined dataset) and for the individual datasets (Drs1 wt/ Drs1 KA, K284A/ Drs1 EQ, E389Q). Classes used for undecimated refinement and model building are highlighted.

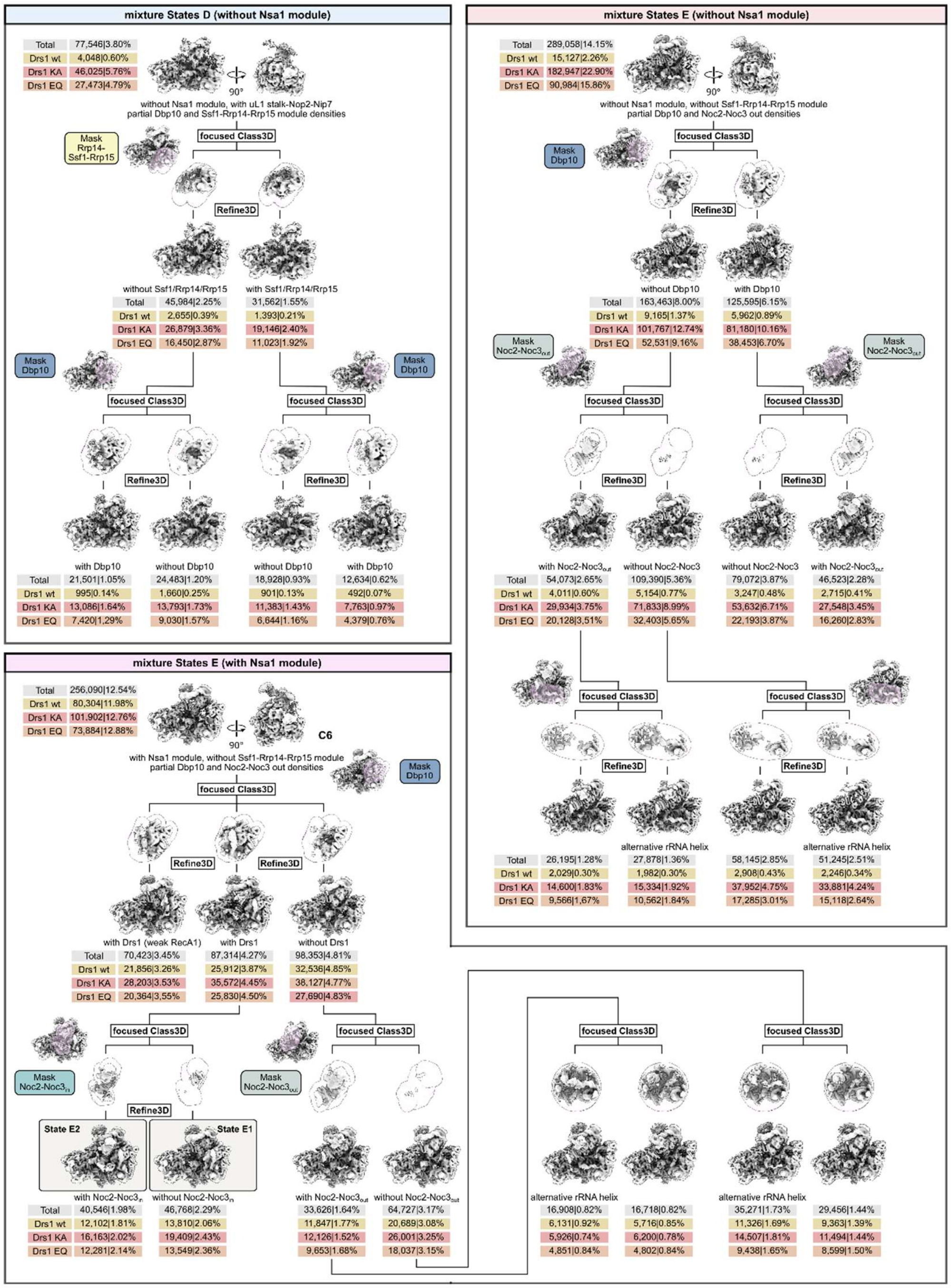

**Supplementary Fig. 6. Processing scheme of the combined Drs1 wt and mutant datasets – Part II.** Sorting schemes of individual classes after the initial round of 3D classification (**Supplementary Fig. 5B**). Particle numbers and their percentages are shown for each class as total (combined dataset) and for the individual datasets (Drs1 wt/ Drs1 KA, K284A/ Drs1 EQ, E389Q). Classes used for undecimated refinement and model building are highlighted.

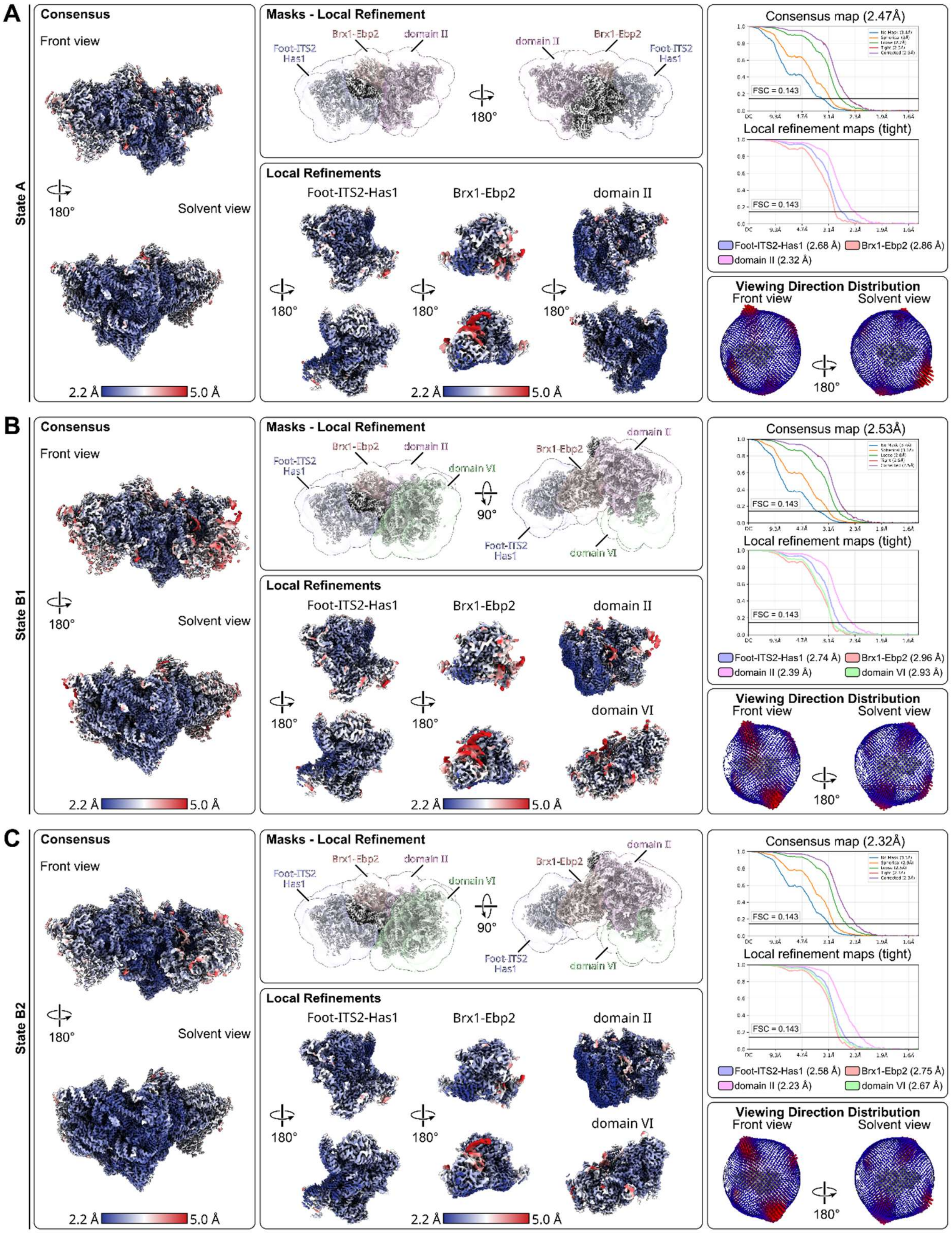

**Supplementary Fig. 7. Cryo-EM validation and local refinements of the combined Drs1 dataset – Part I.** The consensus and local refined maps for States A, B1 and B2 are colored according to their local resolutions. Masks used for the local refinements, Fourier shell correlation (FSC) curves and orientation distributions are shown.

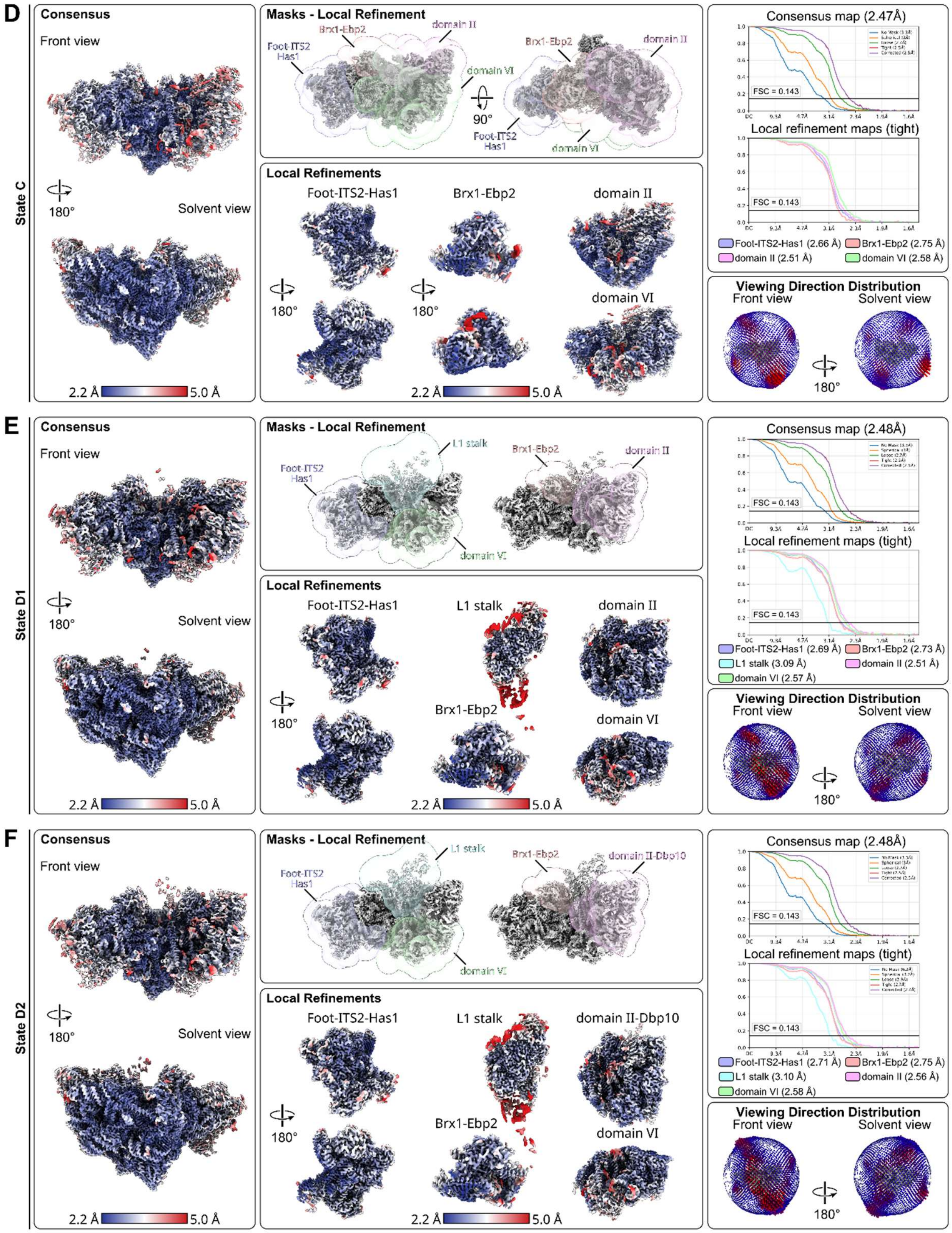

**Supplementary Fig. 8. Cryo-EM validation and local refinements of the combined Drs1 dataset – Part II.** The consensus and local refined maps for States C, D1 and D2 are colored according to their local resolutions. Masks used for the local refinements, Fourier shell correlation (FSC) curves and orientation distributions are shown.

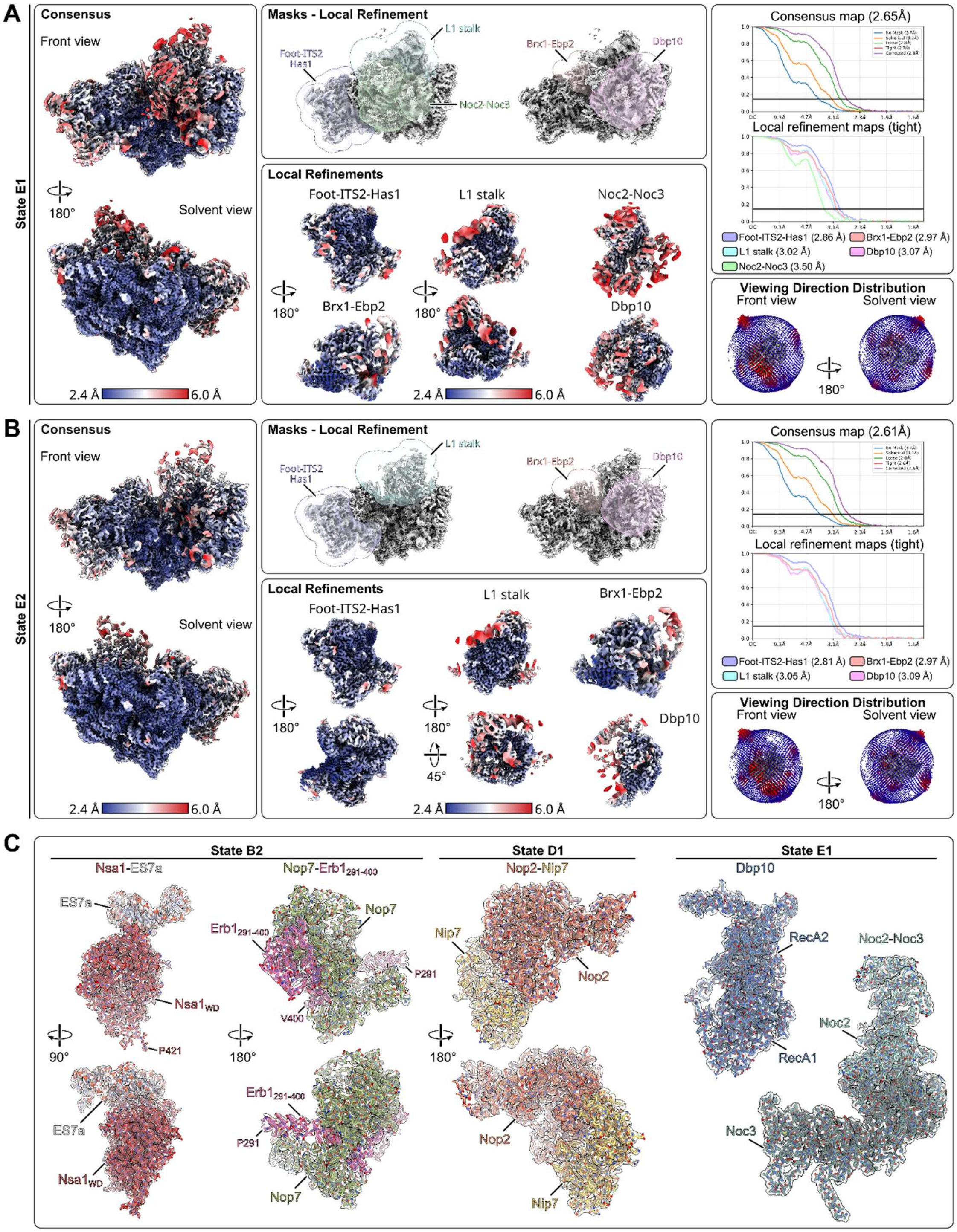

**Supplementary Fig. 9. Cryo-EM validation and local refinements of the combined Drs1 dataset – Part III.** (**A**, **B**) The consensus and local refined maps for States E1 and E2 are colored according to their local resolutions. Masks used for the local refinements, Fourier shell correlation (FSC) curves and orientation distributions are shown. (**C**) Atomic models and the respective segmented cryo-EM densities from State B2, D1 and E1.

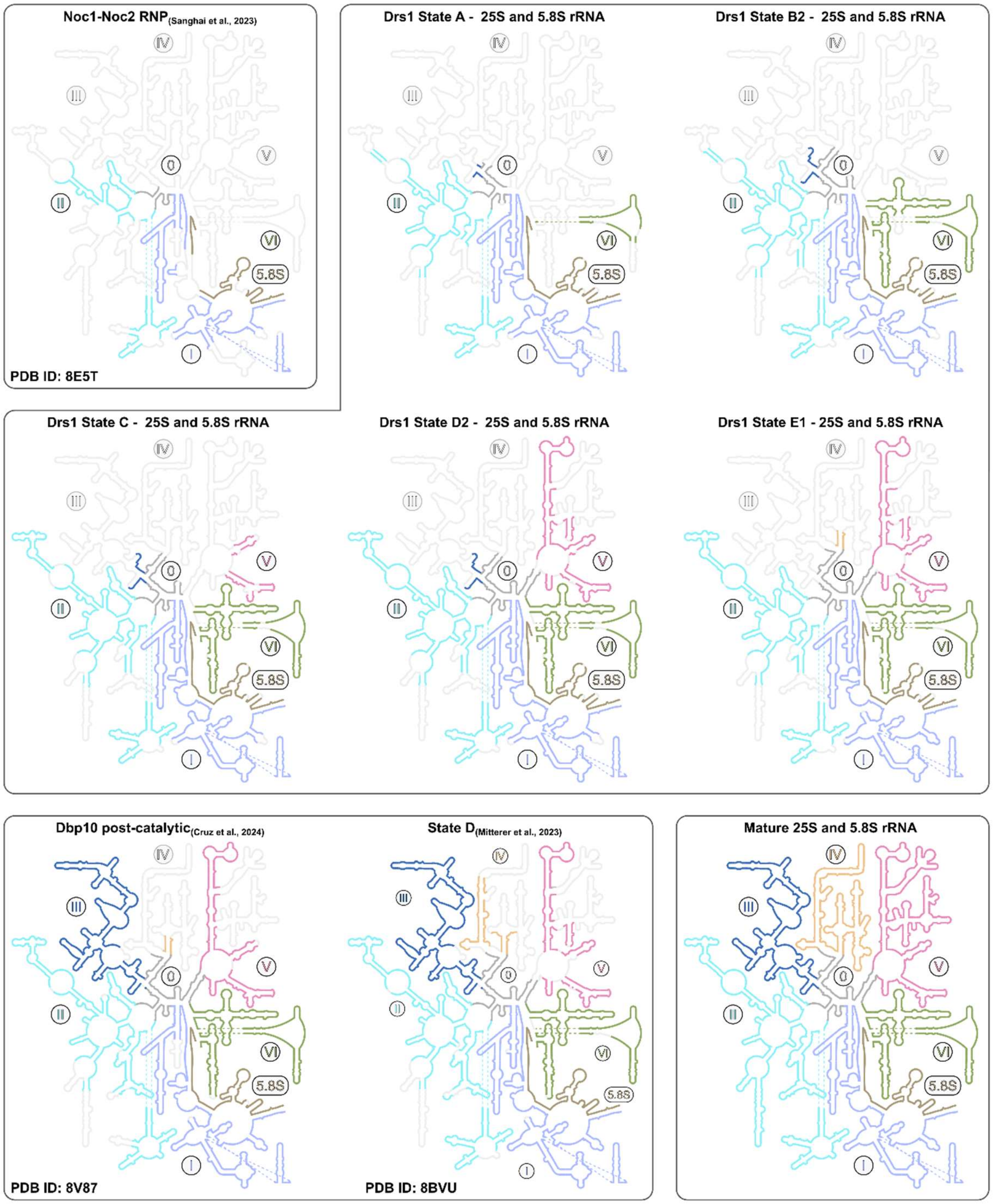

**Supplementary Fig. 10. Modelled rRNA within nucleolar pre-60S states.** Comparisons of incorporated 25S and 5.8S rRNA within the nucleolar pre-60S states purified through Drs1 (middle panels), the preceding Noc1-Noc2 RNP (PDB ID: 8E5T; upper left), as well as subsequent nucleolar pre-60S states with incorporated 25S rRNA domain III (PDB IDs: 8V87 and 8BVU; lower right panels), and the mature 60S subunit (lower left panel) (Cruz et al., 2024; Mitterer et al., 2023; Sanghai et al., 2023). Modelled rRNA regions are colored and labeled and flexible rRNA regions/domains not observed within the individual pre-60S models are shown in transparent gray.

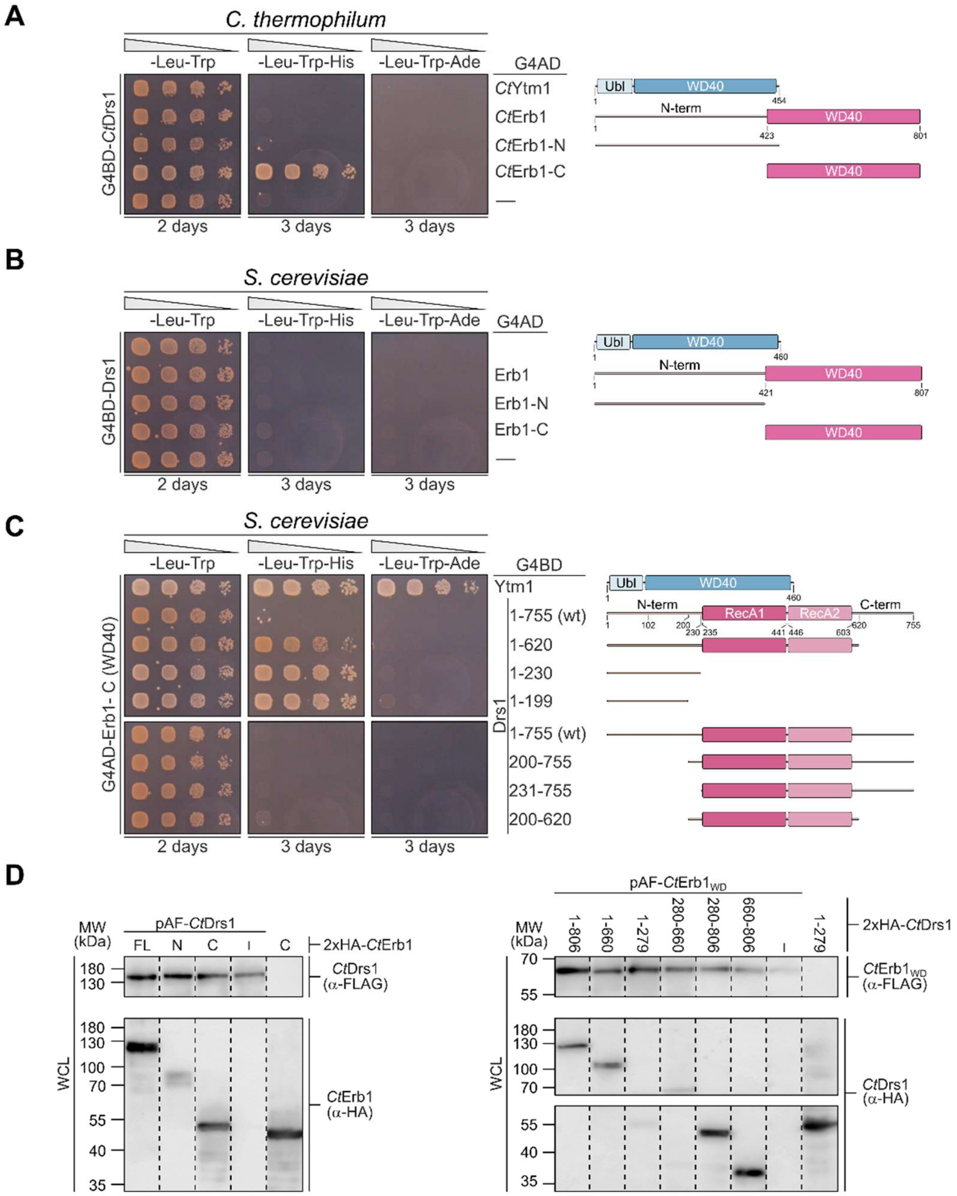

**Supplementary Fig. 11. Protein interaction analysis between Drs1 and Erb1.** (**A-C**) Yeast two-hybrid interaction assays using *C. thermophilum* (A) and *S. cerevisiae* (B-C) constructs. Indicated Drs1, Ytm1 and Erb1 full-length proteins or N- and C-terminal truncation variants were fused to the Gal4 DNA-binding domain (G4BD) or Gal4 activation domain (G4AD), as indicated. Cells were spotted in tenfold serial dilutions on SDC-Leu-Trp (-L-T; plasmid control), SDC-Leu-Trp-His (-L-T-H; growth indicates weak interaction), and SDC-Leu-Trp-Ade (-L-T-A; growth indicates strong interaction) plates and incubated at 30 °C for the indicated times. (**D**) Western blot analysis of the interaction studies shown in **Figure 7C**-**D**. Whole cell lysates (WCL) were analyzed by SDS-PAGE and Western blotting using anti-Flag and anti-HA antibodies to detect the indicated constructs. Membrane images were cut and assembled, as indicated by dotted lines.

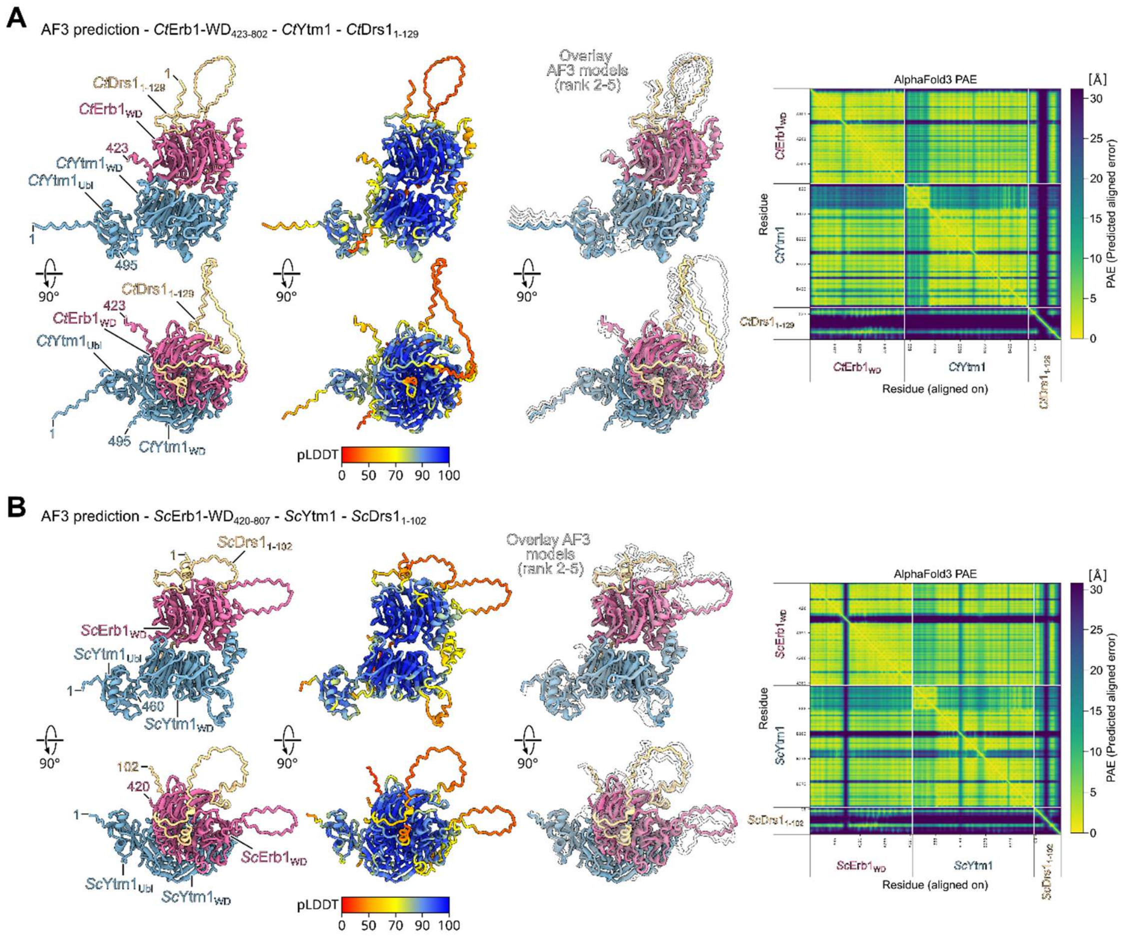

**Supplementary Fig. 12. Alphafold3 predictions of the Drs1-Erb1-Ytm1 interaction.** (**A**, **B**) Alphafold3 (Abramson et al., 2024) predictions of the $Drs1_{N\text{-}term}$-$Erb1_{WD}$-Ytm1 interaction with proteins from *C. thermophilum* (A) and from *S. cerevisiae* (B). The AF3 models are shown labeled and color-coded (left panels), colored according to the pLDDT (middle) and overlayed with the ranked AF3 models 2-5 (middle). The predicted alignment error (PAE) plots are shown (right panels).

**Supplementary Table 1. Cryo-EM data collection, refinement and validation statistics of the combined Drs1 dataset.**

| | pre-60S – State A | pre-60S – State B1 | pre-60S – State B2 | pre-60S – State C |
|---|---|---|---|---|
| **Data collection and processing** | | | | |
| Magnification | 165,000 | 165,000 | 165,000 | 165,000 |
| Voltage (kV) | 300 | 300 | 300 | 300 |
| Electron exposure (e–/Å$^2$) | 40 | 40 | 40 | 40 |
| Defocus range (μm) | 0.5-3.5 | 0.5-3.5 | 0.5-3.5 | 0.5-3.5 |
| Pixel size (Å) | 0.727 | 0.727 | 0.727 | 0.727 |
| Symmetry imposed | C1 | C1 | C1 | C1 |
| Initial particle images (no.) | 2,972,651 | 2,972,651 | 2,972,651 | 2,972,651 |
| Final particle images (no.) | 90,488 | 52,394 | 121,043 | 80,551 |
| Map resolution (Å) | 2.47 | 2.53 | 2.32 | 2.47 |
| FSC threshold | 0.143 | 0.143 | 0.143 | 0.143 |
| Masked map resolution range (Å) | 1.57-38.66 | 1.57-38.62 | 1.92-38.69 | 1.57-38.65 |
| **Refinement** | | | | |
| Model resolution (Å) | 2.3 | 2.4 | 2.2 | 2.4 |
| FSC threshold | 0.5 | 0.5 | 0.5 | 0.5 |
| Map sharpening *B* factor (Å$^2$) | -52 | -50 | -50 | -51 |
| Model composition | | | | |
| Non-hydrogen atoms | 79,701 | 98,045 | 99,885 | 108,704 |
| Protein residues | 6,072 | 7,438 | 7,649 | 8,308 |
| Nucleotide residues | 1,451 | 1,817 | 1,817 | 2,018 |
| Water | 0 | 0 | 0 | 0 |
| Ligands | 1 | 2 | 2 | 2 |
| B factors (Å$^2$) mean | | | | |
| Protein | 38.82 | 39.58 | 32.56 | 37.86 |
| Nucleotide | 50.47 | 54.96 | 43.74 | 52.98 |
| Ligand | 54.50 | 62.20 | 47.47 | 52.60 |
| R.m.s. deviations | | | | |
| Bond lengths (Å) | 0.004 | 0.004 | 0.004 | 0.005 |
| Bond angles (°) | 0.742 | 0.721 | 0.739 | 0.793 |
| Validation | | | | |
| MolProbity score | 1.24 | 1.17 | 1.31 | 1.38 |
| Clash score | 4.02 | 3.82 | 3.90 | 3.97 |
| Poor rotamers (%) | 1.17 | 0.00 | 1.48 | 1.80 |
| Ramachandran plot | | | | |
| Favored (%) | 98.61 | 98.59 | 98.66 | 98.91 |
| Allowed (%) | 1.39 | 1.38 | 1.33 | 1.08 |
| Disallowed (%) | 0.00 | 0.03 | 0.01 | 0.01 |
| Map vs. Model CC (mask) | 0.86 | 0.87 | 0.85 | 0.86 |
| EMDB | | | | |
| Consensus & Local Refinements | EMD-XXXX<br>EMD-XXXX<br>EMD-XXXX<br>EMD-XXXX | EMD-XXXX<br>EMD-XXXX<br>EMD-XXXX<br>EMD-XXXX<br>EMD-XXXX | EMD-XXXX<br>EMD-XXXX<br>EMD-XXXX<br>EMD-XXXX<br>EMD-XXXX | EMD-XXXX<br>EMD-XXXX<br>EMD-XXXX<br>EMD-XXXX<br>EMD-XXXX |
| EMDB (Composite map) | EMD-XXXX | EMD-XXXX | EMD-XXXX | EMD-XXXX |
| PDB | XXXX | XXXX | XXXX | XXXX |

| | pre-60S – State D1 | pre-60S – State D2 | pre-60S – State E1 | pre-60S – State E2 |
|---|---|---|---|---|
| **Data collection and processing** | | | | |
| Magnification | 165,000 | 165,000 | 165,000 | 165,000 |
| Voltage (kV) | 300 | 300 | 300 | 300 |
| Electron exposure (e–/Å$^2$) | 40 | 40 | 40 | 40 |
| Defocus range (μm) | 0.5-3.5 | 0.5-3.5 | 0.5-3.5 | 0.5-3.5 |
| Pixel size (Å) | 0.727 | 0.727 | 0.727 | 0.727 |
| Symmetry imposed | C1 | C1 | C1 | C1 |
| Initial particle images (no.) | 2,972,651 | 2,972,651 | 2,972,651 | 2,972,651 |
| Final particle images (no.) | 83,910 | 74,954 | 40,546 | 46,768 |
| Map resolution (Å) | 2.48 | 2.48 | 2.65 | 2.61 |
| FSC threshold | 0.143 | 0.143 | 0.143 | 0.143 |
| Masked map resolution range (Å) | 1.57-38.61 | 1.57-38.71 | 1.56-42.57 | 1.56-42.54 |
| **Refinement** | | | | |
| Model resolution (Å) | 2.5 | 2.5 | 2.7 | 2.6 |
| FSC threshold | 0.5 | 0.5 | 0.5 | 0.5 |
| Map sharpening *B* factor (Å$^2$) | -51 | -52 | -48 | -49 |
| Model composition | | | | |
| Non-hydrogen atoms | 122,645 | 125,547 | 136,436 | 125,610 |
| Protein residues | 9,410 | 9,800 | 11,156 | 9,857 |
| Nucleotide residues | 2,289 | 2,289 | 2,316 | 2,307 |
| Water | 0 | 0 | 0 | 0 |
| Ligands | 2 | 2 | 2 | 2 |
| B factors (Å$^2$) mean | | | | |
| Protein | 37.34 | 39.36 | 40.41 | 46.53 |
| Nucleotide | 48.66 | 50.41 | 49.66 | 59.61 |
| Ligand | 47.48 | 53.01 | 63.07 | 72.19 |
| R.m.s. deviations | | | | |
| Bond lengths (Å) | 0.005 | 0.004 | 0.004 | 0.006 |
| Bond angles (°) | 0.813 | 0.768 | 0.744 | 0.798 |
| Validation | | | | |
| MolProbity score | 1.38 | 1.35 | 1.23 | 1.20 |
| Clash score | 4.20 | 4.42 | 4.56 | 4.19 |
| Poor rotamers (%) | 1.73 | 1.47 | 0.02 | 0.04 |
| Ramachandran plot | | | | |
| Favored (%) | 98.57 | 98.82 | 98.40 | 98.52 |
| Allowed (%) | 1.41 | 1.17 | 1.58 | 1.47 |
| Disallowed (%) | 0.01 | 0.01 | 0.02 | 0.01 |
| Map vs. Model CC (mask) | 0.87 | 0.86 | 0.85 | 0.86 |
| EMDB | | | | |
| Consensus & Local Refinements | EMD-XXXX | EMD-XXXX | EMD-XXXX | EMD-XXXX |
| | EMD-XXXX | EMD-XXXX | EMD-XXXX | EMD-XXXX |
| | EMD-XXXX | EMD-XXXX | EMD-XXXX | EMD-XXXX |
| | EMD-XXXX | EMD-XXXX | EMD-XXXX | EMD-XXXX |
| | EMD-XXXX | EMD-XXXX | EMD-XXXX | EMD-XXXX |
| | EMD-XXXX | EMD-XXXX | EMD-XXXX | |
| EMDB (Composite map) | EMD-XXXX | EMD-XXXX | EMD-XXXX | EMD-XXXX |
| PDB | XXXX | XXXX | XXXX | XXXX |

**Supplementary Table 2. Yeast strains used in this study.**

| Name | Genotype | Source |
|---|---|---|
| W303 | *ade2-1, his3-11,15, leu2-3,112, trp1-1, ura3-1, can1-100* | (Thomas and Rothstein, 1989) |
| TAPF-*DRS1* | W303 P.*DRS1*-TAP-Flag-*DRS1*::natNT2 | this study |
| pATH-*DRS1* | W303 P.*DRS1*-pA-TEV-(His)$_6$-*DRS1*::natNT2 | this study |
| AID-HA-*DRS1* | W303 P.*DRS1*-AID-HA-*DRS1*::natNT2 P.*ADH1*-Os*TIR1*-9xmyc::*TRP1* | this study |
| AID-HA-*DRS1 NSA1*-FpA | W303 P.*DRS1*-AID-HA-*DRS1*::natNT2 P.*ADH1*-Os*TIR1*-9xmyc::*TRP1 NSA1*-FTpA::HIS3MX6 | this study |
| AID-HA-*DRS1 NOP7*-FpA | W303 P.*DRS1*-AID-HA-*DRS1*::natNT2 P.*ADH1*-Os*TIR1*-9xmyc::*TRP1 NOP7*-FTpA::HIS3MX6 | this study |
| AID-HA-*DRS1 NOP4*-FpA | W303 P.*DRS1*-AID-HA-*DRS1*::natNT2 P.*ADH1*-Os*TIR1*-9xmyc::*TRP1 NOP4*-FTpA::HIS3MX6 | this study |
| AID-HA-*DRS1 UTP10*-FpA | W303 P.*DRS1*-AID-HA-*DRS1*::natNT2 P.*ADH1*-Os*TIR1*-9xmyc::*TRP1 UTP10*-FTpA::HIS3MX6 | this study |
| AID-HA-*DRS1 UTP14*-FpA | W303 P.*DRS1*-AID-HA-*DRS1*::natNT2 P.*ADH1*-Os*TIR1*-9xmyc::*TRP1 UTP14*-FTpA::HIS3MX6 | this study |
| PJ69-4A | *trp1-901, leu2-3,112, ura3-52, his3-200, gal4Δ, gal80Δ, LYS2::GAL1- HIS3, GAL2-ADE2, met2::GAL7-lacZ* | (James et al., 1996) |

**Supplementary Table 3. Plasmids used in this study.**

| Name | Relevant Information | Source |
|---|---|---|
| YEplac181 | *CEN*, *LEU2* | (Gietz and Sugino, 1988) |
| YCplac111-*DRS1* | *CEN, LEU2,* P*DRS1* | this study |
| YCplac111-*drs1*.K284A | *CEN, LEU2,* P*DRS1* | this study |
| YCplac111-*drs1*.D388A | *CEN, LEU2,* P*DRS1* | this study |
| YCplac111-*drs1*.E389Q | *CEN, LEU2,* P*DRS1* | this study |
| YCplac111-TAPF-*DRS1* | *CEN, LEU2,* P*DRS1,* N-terminal TAP-Flag tag | this study |
| YCplac111-TAPF-*drs1*.K284A | *CEN, LEU2,* P*DRS1,* N-terminal TAP-Flag tag | this study |
| YCplac111-TAPF-*drs1*.D388A | *CEN, LEU2,* P*DRS1,* N-terminal TAP-Flag tag | this study |
| YCplac111-TAPF-*drs1*.E389Q | *CEN, LEU2,* P*DRS1,* N-terminal TAP-Flag tag | this study |
| YPGAL111-*DRS1* | *CEN, LEU2,* P*GAL1-10,* | this study |
| YPGAL111-*drs1*.K284A | *CEN, LEU2,* P*GAL1-10,* | this study |
| YPGAL111-*drs1*.D388A | *CEN, LEU2,* P*GAL1-10,* | this study |
| YPGAL111-*drs1*.E389Q | *CEN, LEU2,* P*GAL1-10,* | this study |
| YPGAL111-*drs1*.K284A(aa102-755) | *CEN, LEU2,* P*GAL1-10,* | this study |
| YPGAL111-*drs1*.K284A(aa200-755) | *CEN, LEU2,* P*GAL1-10* | this study |
| YPGAL111-*drs1*.K284A(aa1-679) | *CEN, LEU2,* P*GAL1-10* | this study |
| YPGAL111-*drs1*.K284A(aa1-634) | *CEN, LEU2,* P*GAL1-10* | this study |
| YPGAL111-SV40-NLS-*DRS1* | *CEN, LEU2,* P*GAL1-10*, N-terminal SV40-NLS | this study |
| YPGAL111-SV40-NLS-*drs1*.K284A | *CEN, LEU2,* P*GAL1-10,* N-terminal SV40-NLS | this study |
| YPGAL111-SV40-NLS-*drs1*.K284A(aa102-755) | *CEN, LEU2,* P*GAL1-10,* N-terminal SV40-NLS | this study |
| YPGAL111-SV40-NLS-*drs1*.K284A(aa200-755) | *CEN, LEU2,* P*GAL1-10,* N-terminal SV40-NLS | this study |
| YPGAL111-SV40-NLS-*drs1*.K284A(aa1-679) | *CEN, LEU2,* P*GAL1-10,* N-terminal SV40-NLS | this study |
| YPGAL111-SV40-NLS-*drs1*.K284A(aa1-634) | *CEN, LEU2,* P*GAL1-10,* N-terminal SV40-NLS | this study |
| pG4BDN22-*CtDRS1* | *CEN*, *TRP1*, P*ADH1*, N-terminal Gal4-BD | this study |
| pG4BDN22-*CtDRS1*(aa1-660) | *CEN*, *TRP1*, P*ADH1*, N-terminal Gal4-BD | this study |
| pG4BDN22-*CtDRS1*(aa1-279) | *CEN*, *TRP1*, P*ADH1*, N-terminal Gal4-BD | this study |
| pG4BDN22-*CtDRS1*(aa1-129) | *CEN*, *TRP1*, P*ADH1*, N-terminal Gal4-BD | this study |
| pG4BDN22-*CtDRS1*(aa130-279) | *CEN*, *TRP1*, P*ADH1*, N-terminal Gal4-BD | this study |
| pG4BDN22-*CtDRS1*(aa280-806) | *CEN*, *TRP1*, P*ADH1*, N-terminal Gal4-BD | this study |
| pG4BDN22-*CtDRS1*(aa280-660) | *CEN*, *TRP1*, P*ADH1*, N-terminal Gal4-BD | this study |
| pG4BDN22-*CtDRS1*(aa660-806) | *CEN*, *TRP1*, P*ADH1*, N-terminal Gal4-BD | this study |
| pG4BDN22-*CtYTM1* | *CEN*, *TRP1*, P*ADH1*, N-terminal Gal4-BD | this study |
| pG4ADN111-*CtERB1* | *CEN*, *LEU2*, P*ADH1*, N-terminal Gal4-AD | this study |
| pG4ADN111-*CtERB1*(aa1-454) | *CEN*, *LEU2*, P*ADH1*, N-terminal Gal4-AD | this study |
| pG4ADN111-*CtERB1*(aa423-801) | *CEN*, *LEU2*, P*ADH1*, N-terminal Gal4-AD | this study |
| pG4BDN22-*ScDRS1* | *CEN*, *TRP1*, P*ADH1*, N-terminal Gal4-BD | this study |
| pG4BDN22-*ScDRS1*(aa1-620) | *CEN*, *TRP1*, P*ADH1*, N-terminal Gal4-BD | this study |
| pG4BDN22-*ScDRS1*(aa1-230) | *CEN*, *TRP1*, P*ADH1*, N-terminal Gal4-BD | this study |
| pG4BDN22-*ScDRS1*(aa1-199) | *CEN*, *TRP1*, P*ADH1*, N-terminal Gal4-BD | this study |

| Name | Relevant Information | Source |
|---|---|---|
| pG4BDN22-*ScDRS1*(aa200-755) | *CEN*, *TRP1*, P*ADH1*, N-terminal Gal4-BD | this study |
| pG4BDN22-*ScDRS1*(aa231-755) | *CEN*, *TRP1*, P*ADH1*, N-terminal Gal4-BD | this study |
| pG4BDN22-*ScDRS1*(aa200-620) | *CEN*, *TRP1*, P*ADH1*, N-terminal Gal4-BD | this study |
| pG4BDN22-*ScYTM1* | *CEN*, *TRP1*, P*ADH1*, N-terminal Gal4-BD | (Thoms et al., 2016) |
| pG4ADN111-*ScERB1* | *CEN*, *LEU2*, P*ADH1*, N-terminal Gal4-AD | (Thoms et al., 2016) |
| pG4ADN111-*ScERB1*(aa1-419) | *CEN*, *LEU2*, P*ADH1*, N-terminal Gal4-AD | (Thoms et al., 2016) |
| pG4ADN111-*ScERB1*(aa420-807) | *CEN*, *LEU2*, P*ADH1*, N-terminal Gal4-AD | (Thoms et al., 2016) |
| YEplac181 | 2μ, *LEU2* | (Gietz and Sugino, 1988) |
| YEplac112 | 2μ, *TRP1* | (Gietz and Sugino, 1988) |
| YPGAL181-pATF-*CtDRS1* | 2μ, *LEU2,* P*GAL1-10*, N-terminal protA-TEV-Flag-5xGA tag | this study |
| YPGAL181-pATF-*CtERB1*(aa423-801) | 2μ, *LEU2,* P*GAL1-10*, N-terminal protA-TEV-Flag-5xGA tag | this study |
| YPGAL112-2xHA-*CtERB1* | 2μ, *TRP1,* P*GAL1-10*, N-terminal 2xHA-5xGA tag | this study |
| YPGAL112-2xHA-*CtERB1*(aa1-454) | 2μ, *TRP1,* P*GAL1-10*, N-terminal 2xHA-5xGA tag | this study |
| YPGAL112-2xHA-*CtERB1*(aa423-801) | 2μ, *TRP1,* P*GAL1-10*, N-terminal 2xHA-5xGA tag | this study |
| YPGAL112-2xHA-*CtDRS1* | 2μ, *TRP1,* P*GAL1-10*, N-terminal 2xHA-5xGA tag | this study |
| YPGAL112-2xHA-*CtDRS1*(aa1-660) | 2μ, *TRP1,* P*GAL1-10*, N-terminal 2xHA-5xGA tag | this study |
| YPGAL112-2xHA-*CtDRS1*(aa1-279) | 2μ, *TRP1,* P*GAL1-10*, N-terminal 2xHA-5xGA tag | this study |
| YPGAL112-2xHA-*CtDRS1*(aa280-660) | 2μ, *TRP1,* P*GAL1-10*, N-terminal 2xHA-5xGA tag | this study |
| YPGAL112-2xHA-*CtDRS1*(aa280-806) | 2μ, *TRP1,* P*GAL1-10*, N-terminal 2xHA-5xGA tag | this study |
| YPGAL112-2xHA-*CtDRS1*(aa660-806) | 2μ, *TRP1,* P*GAL1-10*, N-terminal 2xHA-5xGA tag | this study |
| YPGAL112-LEU2D-pA-TEV-*CtYTM1* | 2μ, *TRP1*, *LEU2D*, P*GAL1-10*, N-terminal pA-TEV tag | (Thoms et al., 2016) |
| pET-24d-$(His)_6$-*CtERB1*(aa423-802) | $Kan^r$, T7 promoter, lac operator, N-terminal $(His)_6$ tag | (Thoms et al., 2016) |
| pET-24d-$(His)_6$-*CtDRS1*(aa1-129) | $Kan^r$, T7 promoter, lac operator, N-terminal $(His)_6$-TEV tag | this study |
